# Sensorimotor brain-computer interface performance depends on signal-to-noise ratio but not connectivity of the mu rhythm in a multiverse analysis of longitudinal data

**DOI:** 10.1101/2023.09.30.558407

**Authors:** Nikolai Kapralov, Mina Jamshidi Idaji, Tilman Stephani, Alina Studenova, Carmen Vidaurre, Tomas Ros, Arno Villringer, Vadim Nikulin

**Affiliations:** Department of Neurology, Max Planck Institute for Human Cognitive and Brain Sciences, Leipzig, Germany; International Max Planck Research School NeuroCom, Leipzig, Germany; BIFOLD — Berlin Institute for the Foundations of Learning and Data, Berlin, Germany; Machine Learning Group, Technische Universität Berlin, Berlin, Germany; Max Planck School of Cognition, Leipzig, Germany; Ikerbasque Science Foundation, Bilbao, Spain; TECNALIA, Basque Research and Technology Alliance (BRTA), San Sebastian, Spain; Basque Center on Cognition, Brain and Language, Basque Excellence Research Centre (BERC), San Sebastian, Spain; Department of Neuroscience and Psychiatry, University of Geneva, Geneva, Switzerland; Center for Biomedical Imaging (CIBM), Geneva-Lausanne, Switzerland

## Abstract

**Objective:** Serving as a channel for communication with locked-in patients or control of prostheses, sensorimotor brain-computer interfaces (BCIs) decode imaginary movements from the recorded activity of the user’s brain. However, many individuals remain unable to control the BCI, and the underlying mechanisms are unclear. The user’s BCI performance was previously shown to correlate with the resting-state signal-to-noise ratio (SNR) of the mu rhythm and the phase synchronization (PS) of the mu rhythm between sensorimotor areas. Yet, these predictors of performance were primarily evaluated in a single BCI session, while the longitudinal aspect remains rather uninvestigated. In addition, different analysis pipelines were used to estimate PS in source space, potentially hindering the reproducibility of the results.

**Approach:** To systematically address these issues, we performed an extensive validation of the relationship between pre-stimulus SNR, PS, and session-wise BCI performance using a publicly available dataset of 62 human participants performing up to 11 sessions of BCI training. We performed the analysis in sensor space using the surface Laplacian and in source space by combining 24 processing pipelines in a multiverse analysis. This way, we could investigate how robust the observed effects were to the selection of the pipeline.

**Main results:** Our results show that SNR had both between- and within-subject effects on BCI performance for the majority of the pipelines. In contrast, the effect of PS on BCI performance was less robust to the selection of the pipeline and became non-significant after controlling for SNR.

**Significance:** Taken together, our results demonstrate that changes in neuronal connectivity within the sensorimotor system are not critical for learning to control a BCI, and interventions that increase the SNR of the mu rhythm might lead to improvements in the user’s BCI performance.

## 1. Introduction

A brain-computer interface (BCI) is a system that decodes the intentions of the user based on the recorded activity of their brain and provides commands to external devices (e.g., prostheses; Wolpaw et al. (2002)). These systems have many potential applications ranging from the clinical ones, such as providing a communication pathway for locked-in patients (Chaudhary et al., 2016) or ameliorating symptoms in patients with stroke and Parkinson’s disease (McFarland et al., 2017), to the research ones, such as the detection of mental states and the facilitation of actions in healthy humans (Blankertz et al., 2010b, 2016). Often, BCIs are based on magnetoencephalographic (MEG) or electroencephalographic (EEG) recordings of brain activity. MEG and EEG (M/EEG) have high temporal resolution and provide multiple features of the ongoing or evoked brain activity that can be used as a control signal (Abiri et al., 2019). For example, BCI paradigms based on the P300 component of the evoked response or steady-state visual evoked responses (SSVEP) provide high information transfer rates for efficient communication (Abiri et al., 2019). However, these paradigms always require external stimuli to be presented, which makes the approach less flexible. In contrast, sensorimotor BCIs decode the imaginary movements of limbs or tongue that can be self-initiated and thus provide more flexibility (Leeb et al., 2007; Yuan and He, 2014; Scherer and Vidaurre, 2018). Decoding of the imaginary movements is often based on the modulation of power in the alpha (8 – 13 Hz) and beta (13 – 30 Hz) frequency ranges in sensorimotor brain areas, also referred to as event-related desynchronization or synchronization (ERD/ERS; Pfurtscheller and Lopes da Silva (1999); Pfurtscheller et al. (1996)). Sensorimotor BCIs are also used to facilitate the recovery of motor functions during rehabilitation after a stroke (Cervera et al., 2018; Kruse et al., 2020; Peng et al., 2022).

While BCI seems to be a promising approach with multiple clinical applications, some participants remain unable to control it (Allison and Neuper, 2010). Typically, participants complete several training sessions to learn to control a BCI. However, their performance in the task varies considerably, and on average around 20% of the participants fail to learn the task (Sannelli et al., 2019). The mechanisms underlying successful modulation of brain activity for controlling a BCI are not clear yet. However, previous studies have identified several psychological (Hammer et al., 2012; Jeunet et al., 2015) and neurophysiological (Blankertz et al., 2010a; Sugata et al., 2014; Samek et al., 2016; Vidaurre et al., 2020; Jorajuría et al., 2023) predictors of successful control of a sensorimotor BCI. These predictors allow pre-screening of participants to provide the full training only if the participant is likely to successfully control the BCI (Sannelli et al., 2019).

Neurophysiological predictors of successful BCI control also provide information about the features of brain activity (e.g., neuronal networks) that play a role in the success of BCI training. For example, the signal-to-noise ratio (SNR) of the sensorimotor mu rhythm during resting state was positively correlated (*r* = 0.53) with the online accuracy of sensorimotor BCI control (Blankertz et al., 2010a). The SNR was defined as the maximal ratio of the periodic and aperiodic (1/f noise) components of the power spectrum in the 2-35 Hz frequency range. This predictor was later validated in an independent dataset with a similar experimental paradigm (Acqualagna et al., 2016). Moreover, several other neural correlates of performance in a sensorimotor BCI task are related to the SNR of the mu rhythm, for example, the performance potential factor (Ahn et al., 2013) or the spectral entropy at C3 electrode during resting-state (Zhang et al., 2015).

Although SNR seems to be a well-established predictor of BCI performance, it is often investigated in the context of a single BCI session. However, the relationship between SNR and performance could change if participants with low SNR eventually learned the task or if the SNR changed throughout a multi-session BCI training. Therefore, it is crucial to validate this predictor in a longitudinal analysis, which is one of the aims of the current study.

Other predictors of sensorimotor BCI performance include long-range temporal correlations (Samek et al., 2016), functional connectivity between sensorimotor brain regions (Sugata et al., 2014; Vidaurre et al., 2020), and the strength of mu vs. beta phase-phase coupling (Jorajuría et al., 2023). Connectivity-based predictors might be especially relevant since motor imagery involves activation of multiple interacting brain areas (Solodkin et al., 2004; Halder et al., 2011; Hardwick et al., 2018). The strength and the phase lag of these interactions can be quantified using various connectivity measures and then related to the performance in the sensorimotor BCI task. Thereby, connectivity could provide additional information about the underlying neuronal networks that is not reflected in the SNR.

When considering M/EEG-based functional connectivity within the same (e.g., alpha/mu) frequency band, phase synchronization (PS) and amplitude envelope correlation (AEC) can reflect different properties of the underlying neuronal networks. Studies combining EEG and fMRI (functional Magnetic Resonance Imaging) have previously shown that the power of alpha and beta oscillations at C3 and C4 is negatively correlated with the blood-oxygen-level-dependent (BOLD) fMRI signal in sensorimotor areas during the execution of real and imaginary hand movements (Ritter et al., 2009; Yuan et al., 2010). Therefore, AEC primarily captures the low-frequency (below 0.1 Hz) dynamics of brain activity similar to the fMRI connectivity based on the BOLD signal (Engel et al., 2013). In contrast, phase synchronization between high-frequency (above 5 Hz) oscillations might reveal additional information that is only accessible with the high temporal resolution of M/EEG (Engel et al., 2013). In particular, phase synchronization was proposed to be a mechanism of efficient communication between neuronal populations (Engel et al., 2001; Fries, 2005; Palva and Palva, 2007) and can reflect short-term changes in the functional organization of neuronal networks due to plasticity (Engel et al., 2013). Therefore, in the current study, we also investigated the role of phase synchronization of the sensorimotor mu (9-15 Hz) oscillations in the successful control of a sensorimotor BCI.

Several studies have already applied various M/EEG-based phase synchronization measures in the context of sensorimotor BCI training. First, BCI performance was positively correlated with the imaginary part of coherency (ImCoh; Nolte et al. (2004)) of the mu rhythm between sensorimotor areas both before and during the trial (Sugata et al., 2014; Vidaurre et al., 2020). In addition, the phase locking value (PLV; Lachaux et al. (1999)) of alpha-band oscillations within the motor areas of the right hemisphere was higher for the successful participants in comparison to the unsuccessful ones (Leeuwis et al., 2021). Finally, in a whole-head analysis, Corsi et al. (2020) observed a global decrease in ImCoh during motor imagery compared to resting state. While phase synchronization seems to play a role in sensorimotor BCI training, the results were obtained using various PS measures and partially in the context of single-session experiments. To address these issues, we examined several PS measures and ran a longitudinal analysis of changes in phase synchronization and its relationship with the BCI performance.

Studies investigating longitudinal changes in phase synchronization are scarce in the sensorimotor BCI literature. On the one hand, Corsi et al. (2020) observed a progressive decrease of ImCoh during motor imagery in alpha and beta bands along sessions. On the other hand, the positive correlation between ImCoh and BCI performance in one session (Sugata et al., 2014; Vidaurre et al., 2020) may suggest the entrainment of task-relevant networks throughout the training. However, in both cases, ImCoh reflects a mixture of the strength and the phase lag of the interaction between brain areas, which can only be disentangled with other PS measures, such as coherence. Therefore, further validation of these results in the longitudinal setting with multiple PS measures is necessary.

In practice, the estimation of phase synchronization in M/EEG critically depends on the proper control for confounding factors (Bastos and Schoffelen, 2015). In the current study, we focused on the effects of volume conduction and signal-to-noise ratio (SNR). To overcome these challenges, we used PS measures, which are insensitive to zero-lag interactions (e.g., ImCoh), and applied a correction for SNR in the statistical analysis.

Furthermore, to obtain a higher spatial specificity of the estimated PS values, we performed the source space analysis using time courses of brain activity in particular regions of interest (ROIs). For this purpose, two-step processing pipelines are typically used (Schoffelen and Gross, 2009). First, inverse modeling is applied to reconstruct time courses of activity for individual sources within the cortex. Second, time courses of activity for all sources within the ROI are aggregated to extract one or several time courses of activity in the ROI. While multiple approaches exist for inverse modeling and extraction of ROI time series, there is no consensus on the most appropriate pipeline in the community. Previous studies have shown that the choice of methods for inverse modeling and extraction of ROI time series affects the estimated PS values in real and simulated data (Mahjoory et al., 2017; Pellegrini et al., 2023). Therefore, multiple pipelines should be considered simultaneously to arrive at a valid conclusion about genuine neuronal connectivity based on M/EEG data.

To address the multitude of possible pipelines while analyzing SNR and phase synchronization as predictors of BCI performance, we ran a multiverse analysis (Steegen et al., 2016) using several pipelines for extraction of ROI time series. While results are typically reported only for one or a few of many possible pipelines, the idea of the multiverse analysis is to consider a set of reasonable pipelines and report the results for all of the considered options. This way, one can not only analyze the variability of the estimated PS values similar to Mahjoory et al. (2017) but also assess the robustness of the observed effects (e.g., on BCI performance) to the selection of the pipeline. More pronounced effects should be more robust to changes in the processing pipeline, and including several pipelines in the analysis may reveal important information about the influence of different processing steps on the observed results.

Overall, in the current study, we aimed to validate and extend the findings about the effects of SNR and PS of the mu rhythm on BCI performance in a publicly available longitudinal dataset (Stieger et al., 2021). We focused on four sensorimotor ROIs corresponding to the primary motor and somatosensory cortices. These ROIs were previously shown to be the most involved in the BCI training based on imaginary movements (Samek et al., 2016; Vidaurre et al., 2020; Nierhaus et al., 2021). In the current analysis, we aimed to address the following research questions:

1. Do SNR and PS predict performance not just in one but also in multiple training sessions?
2. Do SNR and PS change over time due to BCI training?
3. Are SNR, PS, and the observed effects for questions 1 and 2 robust to the selection of processing steps in the source space analysis?

To touch upon the open questions regarding the multitude of existing approaches for source space analysis and estimation of phase synchronization, we considered a set of existing methods and performed a multiverse analysis to capture the between-pipeline variability in estimated values of SNR and PS, their effects on BCI performance, and longitudinal changes over time. In addition, to ensure the end-to-end repeatability of the results, we designed the analysis pipeline to automatically include the results in a publishable report, which, as we hope, will be useful as a template for future studies involving a multitude of different analysis pipelines.

## 2. Materials and Methods

### 2.1. Description of the Dataset

We used publicly available EEG recordings of 62 participants (50 female; 55 right-handed; mean age = 39.2 years, SD = 14.1 years) from a study that investigated the effects of mindfulness-based training on performance in a motor imagery BCI task (Stieger et al., 2021). Participants first completed a baseline BCI training session and then were randomly assigned to an 8-week mindfulness intervention (n = 33; 26 female; 28 right-handed; mean age = 42.2, SD = 14.7) and a wait-list control condition of the same length (n = 29; 24 female; 27 right-handed; mean age = 35.8, SD = 12.9). After eight weeks, participants returned to the lab for 6-10 more sessions of BCI training (Fig. 1A). All experiments were approved by the institutional review boards of the University of Minnesota and Carnegie Mellon University. Informed consent was obtained from all participants.

**Figure 1:**
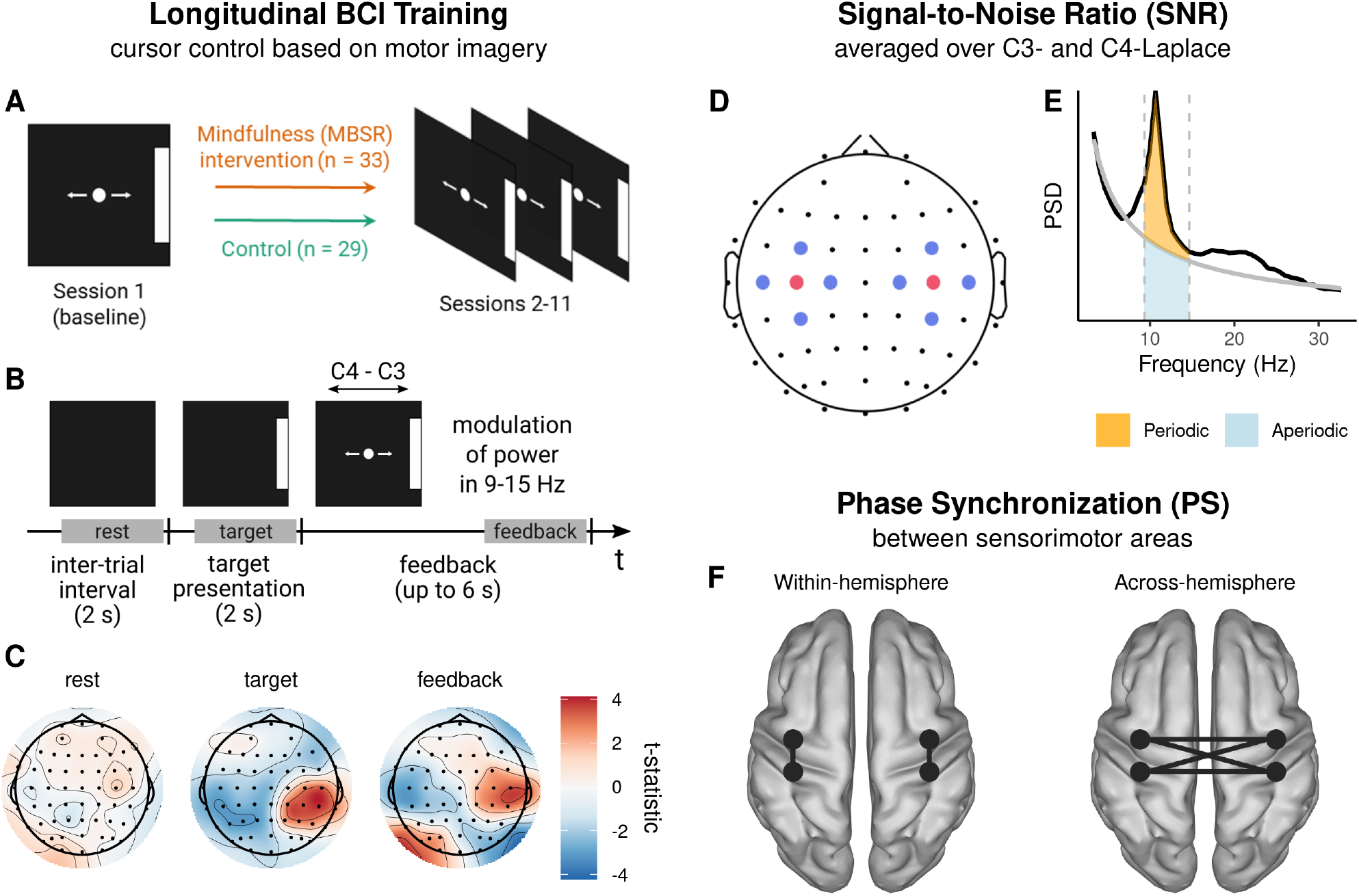
A publicly available dataset (Stieger et al., 2021) was used to test two previously described predictors of successful BCI performance (Blankertz et al., 2010a; Vidaurre et al., 2020) in the longitudinal setting with multiple pipelines for source space analysis. (A) Participants were assigned to MBSR (mindfulness-based stress reduction) and control groups and completed up to 11 sessions of cursor control training. (B) Trial structure of the horizontal cursor control task with time windows of interest highlighted (rest: [0.49, 1.99] s of the inter-trial interval, target: [0.49, 1.99] s of the target presentation interval, feedback: [-1.51, -0.01] s relative to the end of the feedback interval). Participants performed imaginary movements of their left and right hands to control a cursor, whose position was calculated based on the values of mu power at Laplace-filtered channels C3 and C4 in real time. Note that the illustrations of the provided feedback in Panels A and B are schematic and might not exactly reflect what participants saw on the screen. (C) Sensor space channel-wise t-statistic of difference in mu power between trials that involved imaginary movements of the right and left hand. While no difference in mu power was observed during the resting-state period, effects emerged over sensorimotor areas during target presentation, accompanied by effects over visual areas due to the movement of the cursor during the feedback period. (D) Surface Laplacian was applied in the offline analysis to channels C3 and C4 (red) using the neighboring channels (blue) before estimating SNR. (E) SNR was estimated as the ratio of the total (periodic + aperiodic) power and the power of the aperiodic component in the 9-15 Hz frequency range. The gray line depicts the 1/f fit obtained with FOOOF. (F) Phase synchronization (PS) between sensorimotor ROIs was estimated in source space, and PS values were averaged over the highlighted within-hemisphere and across-hemisphere connections.

### 2.2. Experimental Procedure

During each BCI session, participants performed imaginary movements (opening and closing) of their hands to control a cursor, which was displayed on the screen in front of them in the BCI2000 system (Schalk et al., 2004). Each session included three tasks: (1) horizontal cursor control task (via imaginary movements of the left or right hand), (2) vertical cursor control task (down: voluntary rest, up: imaginary movement of both hands), (3) 2D control task (the targets were reachable through one of the previous strategies, but the cursor moved in both directions). Each task included 150 trials, and the number of trials was balanced across classes for both 1D and 2D control tasks. In the current study, we only analyzed the EEG data from the first (horizontal cursor control) task since it was shown to be more tightly related to sensorimotor areas than the second task (Stieger et al., 2020). At the same time, the participants’ performance was similar across all three tasks (Supplementary Material, Section A), so the main qualitative findings of the analyses should also apply to other tasks.

The structure of all trials is shown in Fig. 1B. First, participants saw a blank screen during the inter-trial interval of 2 s. Then, a bar appeared on one of the sides of the screen, indicating the target action to execute. After 2 seconds of target presentation, a cursor (circle) appeared in the middle of the screen, and its position was calculated based on the EEG data acquired in real time. Trials ended either when the cursor reached any side of the screen (not necessarily the target one) or after the timeout when 6 seconds passed without any target being reached.

Feedback was presented with a cursor, whose position was updated in real time based on the EEG power in the mu (9-15 Hz) frequency range. Power was calculated based on an autoregressive model of order 16 fitted to the most recent 160 ms of the EEG data after applying the surface Laplacian to channels C3 and C4 (using the closest neighboring channels FC3, CP3, C1, C5 and FC4, CP4, C2, C6, respectively). The horizontal position of the cursor was determined by the lateralization of mu power (C4 – C3), while the vertical position reflected the total mu power (C4 + C3). Feedback values were re-calculated every 40 ms and normalized by subtracting the mean and dividing over the standard deviation. The mean and the standard deviation were constantly updated based on the last 30 seconds of data. More details about the experimental procedure can be found in (Stieger et al., 2020, 2021).

### 2.3. EEG Acquisition

EEG was acquired using SynAmps RT amplifiers and Neuroscan acquisition software (Compumedics Neuroscan, VA). Data were recorded with a sampling frequency of 1 kHz and band-pass filtered between 0.1 and 200 Hz with an additional notch filter at 60 Hz. EEG data were acquired from 62 channels with the following locations according to the 10-5 system: Fp1, Fpz, Fp2, AF3, AF4, F7, F5, F3, F1, Fz, F2, F4, F6, F8, FT7, FC5, FC3, FC1, FCz, FC2, FC4, FC6, FT8, T7, C5, C3, C1, Cz, C2, C4, C6, T8, TP7, CP5, CP3, CP1, CPz, CP2, CP4, CP6, TP8, P7, P5, P3, P1, Pz, P2, P4, P6, P8, PO7, PO5, PO3, POz, PO4, PO6, PO8, CB1, O1, Oz, O2, CB2. AFz was used as the ground electrode, while the reference electrode was between Cz and CPz.

### 2.4. Preprocessing

EEG preprocessing and analyses were performed in MATLAB R2022b (The MathWorks; RRID: SCR_001622) using custom scripts employing functions from EEGLAB 2021.0 (Delorme and Makeig (2004); RRID: SCR_007292), BBCI (Blankertz et al., 2016), Brainstorm (Tadel et al. (2011); RRID: SCR_001761), MVGC (Barnett and Seth (2014); RRID: SCR_015755) and METH (Guido Nolte; RRID: SCR_016104) toolboxes. For source space visualizations, we utilized functions from (Haufe and Ewald, 2019).

First, trials were concatenated to restore continuous segments of data accounting for breaks during the recording. Then, EEG time series were downsampled to 250 Hz, and channels CB1 and CB2 were removed as they are not part of the 10-10 system. A semi-automatic identification of bad trials, channels, and components was applied as follows. Trials and channels were rejected if the z-score of power within 1-45 Hz was higher than three in at least 5% of trials for a certain channel or in at least 5% of channels for a certain trial. This procedure was performed recursively until nothing could be rejected. Additionally, we used the clean_rawdata EEGLAB plugin to reject channels if one of the following conditions was met: (1) the variance of the channel data was near zero for at least five seconds, (2) the ratio of the power of the line noise and power of the signal below 50 Hz exceeded 4, or (3) the correlation of the channel data with an interpolated estimate based on the data from neighboring channels was less than 0.8. After the removal of bad trials and channels, EEG data were re-referenced to the common average reference and filtered with a forward-backward second-order high-pass Butterworth filter with a cutoff frequency of 1 Hz. Then, we applied independent component analysis (ICA) based on the FastICA approach (Hyvärinen, 1999) and used ICLabel (Pion-Tonachini et al., 2019) for distinguishing ICA components of different types: brain, muscle, eye, heart, line noise, and channel noise. Based on the output of ICLabel, components that explained 95% of the variance in the data were rejected if their probability of originating from the brain was less than 20%, and other components were rejected only if their probability of belonging to one of the non-brain classes was at least 80%.

Results of the automatic preprocessing were verified through visual inspection of power spectra in sensor space as well as topographic maps and power spectra of kept and rejected ICA components. Overall, 3 sessions were excluded from the analysis due to poor data quality. Then, we removed previously identified bad trials, channels, and ICA components from the raw EEG data that were not high-pass filtered. The removed channels were interpolated, and EEG time series were downsampled to 250 Hz. DC offset was removed by subtracting the mean of the signal within continuous data segments. The resulting data were used for the analyses described below.

### 2.5. Overview of the Analyses

In this subsection, we provide a brief overview of the performed analyses. A detailed description of the processing steps is presented in the subsequent subsections.

In the current study, we only analyzed the data from the first (horizontal cursor control) task, which was based on the imaginary movements of the left or right hand. Additionally, we combined the data from both participant groups since a previous analysis of the same dataset has shown that the mindfulness intervention did not affect the performance in the horizontal cursor control task (Stieger et al., 2020).

We estimated the values of SNR of the mu rhythm and phase synchronization (PS) between sensorimotor areas to investigate their relationship with BCI performance and changes over time. For both analyses, we focused on the oscillations in the 9-15 Hz frequency range and the [0.49, 1.99] s window of the inter-trial interval (labeled as rest in Fig. 1B). During this interval, participants did not perform any task similar to a typical resting-state recording, and previous studies often used resting-state data to predict BCI performance in subsequent training sessions. Additionally, we considered the [0.49, 1.99] s window of the target presentation interval (target, Fig. 1B) as well as the [-1.51, -0.01] s window relative to the end of the feedback interval (feedback, Fig. 1B) for the analysis of task-related changes in mu power.

The performance in the BCI task (online accuracy) was assessed with the percentage of correct trials among those that did not end due to timeout. Trials were considered correct if the cursor reached the target side of the screen. To make the results of the study applicable to commonly used decoding-based BCI paradigms, we also considered offline accuracy and area under the receiver operating characteristic curve (AUC) as alternative metrics for the evaluation of performance (Supplementary Material, Section B). Since the values of online accuracy and offline performance metrics showed high between- and within-subject correlations (Fig. S2, Tab. S2), we performed the main analysis only for the online accuracy (later referred to as accuracy).

#### Sensor space analysis

As a sanity check and a replication of results by Stieger et al. (2020), we first computed the difference in mu (9-15 Hz) power in the aforementioned time windows of interest to ensure that task-related contrasts can be observed during target presentation and feedback but not the inter-trial interval (Fig. 1C).

#### Laplacian-based analysis

Before estimating the SNR of the mu rhythm, we applied the surface Laplacian by subtracting the mean of the neighboring channels (FC3, C5, C1, CP3 or FC4, C6, C2, CP4) from the data at channels C3 and C4 (Fig. 1D). The same transformation was used during the experiment for calculating the feedback values in real time. We estimated the SNR of the mu rhythm (Fig. 1E) and correlated it with the BCI performance similar to (Blankertz et al., 2010a). Additionally, we examined longitudinal changes in SNR across sessions to find out whether the BCI training affected the SNR of the mu rhythm. Finally, we also estimated values of PS between Laplace-filtered channels C3, C4, CP3, and CP4, which are located directly over the sensorimotor areas. The results of the Laplacian-based analysis of PS are described in the Section C of the Supplementary Material as the conclusions were very similar to the source space analysis described below.

#### Source space analysis

To extend the results of (Vidaurre et al., 2020) to the longitudinal setting, we estimated the SNR of the mu rhythm and PS between the sensorimotor regions of interest (ROIs) in source space. The time courses of activity in each ROI were computed through inverse modeling and subsequent aggregation of reconstructed time series of source dipoles within the ROI. Various methods for inverse modeling and extraction of ROI time series are used in the literature with few guidelines for preferring one over the other. Therefore, we combined several widely used data-driven and data-independent approaches in a multiverse analysis (Steegen et al., 2016) to investigate the robustness of SNR and PS values as well as related statistical effects (e.g., on BCI performance) to the selection of the pipeline (Fig. 2A).

**Figure 2:**
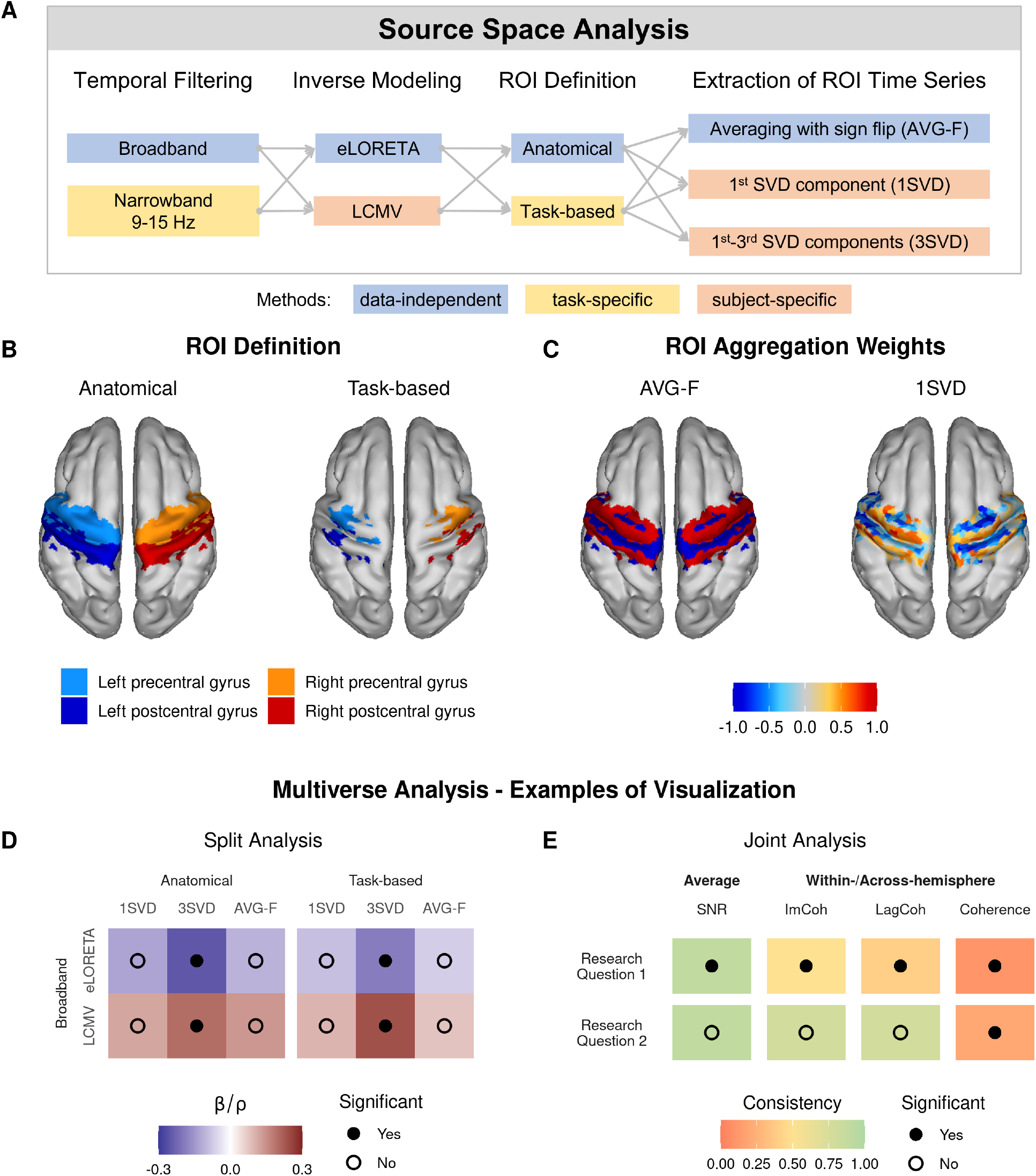
Overview of the multiverse analysis of SNR and phase synchronization in the source space. (A) 24 combinations of the data-independent and task- or subject-specific methods were used in the current analysis. (B) Anatomical and task-based (derived using CSP) definitions of sensorimotor ROIs. (C) AVG-F and 1SVD weights for all sources within sensorimotor ROIs for an exemplary subject. (D) In the split multiverse analysis, statistical results were aggregated in a table to assess the robustness of effects to the selection of the pipeline. Estimated correlations (*ρ*, between-subject effect) or regression coefficients (*β*, within-subject effect) were coded with color, and the significance of the effects was indicated by filled black dots. (E) In the joint multiverse analysis, data from all pipelines were pooled to obtain one result for each performance predictor and research question. Color codes consistency – the number of individual pipelines that led to the same result in terms of significance and, if significant, direction of the effect as the joint analysis.

### 2.6. Forward Model

We used the “New York Head” forward model (Huang et al., 2016), which was derived using the finite element method based on the ICBM152 anatomical template (Fonov et al., 2009, 2011). The model contains several lead field matrices calculated for different numbers and orientations of the source dipoles (later referred to as sources). We used the lead field matrix for 4502 sources with fixed orientations perpendicular to the cortical surface. Since channels PO5 and PO6 were not included in the precomputed lead field, we excluded them before source space analysis. The common average reference transform was applied to the lead field matrix to match the preprocessing of the EEG data.

### 2.7. Inverse Modeling

We used two inverse solutions with different underlying assumptions: eLORETA (Pascual-Marqui, 2007) and linearly constrained minimal variance (LCMV) beamformer (Van Veen et al., 1997). For both approaches, we used the implementation from the METH toolbox (Guido Nolte; RRID: SCR_016104) with minor modifications from (Haufe and Ewald, 2019). The regularization parameter was set to 0.05 and the identity matrix was used as the noise covariance matrix.

eLORETA is a data-independent approach that belongs to the family of weighted minimum norm inverse solutions and provides zero source localization error (Pascual-Marqui et al., 2011). In the described setting, this approach is also data-independent. In contrast, LCMV is a data-driven method and is fit to the covariance matrix of the data. We averaged covariance matrices for both imaginary movements and calculated a separate LCMV beamformer for each subject and session.

### 2.8. Extraction of ROI Time Series

After the inverse modeling, one obtains a reconstructed time series of activity for each source. Considering the spatial resolution of EEG, it is reasonable to reduce the dimensionality of the source space. The common approach is to aggregate time courses of activity of sources within each ROI into a single or several time series. Yet, multiple aggregation methods exist in the literature, and there is no consensus in the community on the most appropriate method. In particular, previous studies have used averaging (Babiloni et al., 2005), averaging with sign flip (AVG-F; Lai et al. (2018)), singular value decomposition (SVD; Rubega et al. (2019)), etc. In the current analysis, we considered AVG-F and SVD to compare commonly used data-independent and data-driven approaches.

For both approaches, the time series of activity for all sources within the ROI are concatenated to form a matrix. By fitting SVD, one decomposes the multivariate time series of activity into components sorted by the explained variance of the reconstructed source data. Then, a few first components are selected to represent the activity of the whole ROI. We considered either only the first (1SVD) or the first three components (3SVD) as performed in, e.g., (Rubega et al., 2019; Pellegrini et al., 2023) or (Vidaurre et al., 2020; Pellegrini et al., 2023), respectively.

Alternatively, AVG-F assigns equal weights to all sources within the ROI, and a sign flip is applied to some sources to prevent the cancellation of the activity of dipoles with opposite orientations. The sign flip is especially important for the current study since a forward model with fixed dipole orientations was used. To determine the sources that should be flipped, SVD is applied to the leadfield of sources within the ROI to find the dominant orientation of source dipoles. If the angle between the orientation of the dipole and the dominant orientation is larger than 90 degrees, the time series corresponding to this dipole is flipped (that is, multiplied by a negative one). We used the implementation of sign flip from Brainstorm (Tadel et al., 2011). Fig. 2C shows 1SVD and AVG-F weights for all sources within the sensorimotor ROIs based on the data of an exemplary subject.

### 2.9. Anatomical and Task-Based Definitions of ROIs

All the analyses in the source space were performed for four sensorimotor ROIs — pre- and postcentral gyri of both hemispheres — either according to their definitions in the Harvard-Oxford atlas (Frazier et al., 2005; Desikan et al., 2006; Makris et al., 2006; Goldstein et al., 2007; Jenkinson et al., 2012) or reduced to a group of task-relevant sources (Fig. 2B). To select a subset of sources that contribute the most to the observed task-related changes in brain activity, we applied a mask in source space derived from the common spatial pattern (CSP) transformation (Koles et al., 1990; Ramoser et al., 2000). CSP was applied to the sensor space data filtered in the 9-15 Hz range for extracting spatial filters that explain the most difference in EEG power between the two imaginary movements. For this purpose, we used the EEG data during the [0.49, 1.99] s window of the target presentation interval. Covariance matrices of the signal were calculated for each subject, session, and imaginary movement separately. Then, for each subject and session, covariance matrices corresponding to different imaginary movements were normalized to make the trace of their average equal to one. The normalization allowed us to exclude the difference in signal power between subjects and sessions while preserving the within-session difference in power between channels and imaginary movements. Normalized covariance matrices were averaged over all subjects and sessions and then used to obtain one set of CSP filters and patterns for all participants. CSP patterns were then source reconstructed with eLORETA. A threshold based on the 97.5th percentile of activity strength was applied to select the most responsive sources, which formed the resulting source mask. The mask was applied to the anatomical definitions of sensorimotor ROIs to obtain a task-based reduced representation.

### 2.10. Filtering

Due to the 1/f shape of the M/EEG power spectra, lower frequencies (< 7 Hz) might have higher power and overshadow mu oscillations in covariance calculations (Chalas et al., 2022). By filtering the data in a narrow frequency band, one makes sure that data-dependent methods (LCMV, SVD) are not affected by frequencies outside of the target band. At the same time, data-independent methods (eLORETA, AVG-F) are not affected by filtering.

To investigate how filtering in a narrow frequency band affects data-dependent methods for inverse modeling and extraction of ROI time series, we considered two cases: broadband with no filtering (BB) or band-pass filtering in the 9-15 Hz band (NB). A forward-backward fourth-order Butterworth filter was applied before restricting the data to the time windows of interest and applying the inverse modeling. Since the recording contained breaks, 8 seconds of data at the beginning and the end of continuous data segments were mirrored to minimize filtering-related edge effects. Separate LCMV beamformers and sets of SVD weights were calculated for broadband and narrowband data. The computed weights were applied to the broadband data before estimating SNR and PS.

### 2.11. Signal-to-Noise Ratio

Signal-to-noise ratio (SNR) was estimated as the ratio of the total power and the power of the aperiodic component of the signal in the 9-15 Hz frequency range:

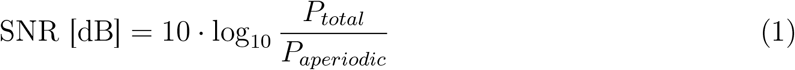

The aperiodic component of the signal was estimated using FOOOF (Fig. 1E; Donoghue et al. (2020)) with the following set of parameters: 1-45 Hz fit range, 2-12 Hz as limits of peak width, and 3 as the maximal number of peaks. In the case of very low mu power, negative values may be obtained for the power of the periodic component, and total power was used in the definition of the SNR to prevent computational errors when taking the logarithm of the power ratio. Values of SNR were estimated in the same manner for the Laplace-filtered sensor space data and in the source space, later referred to as Laplace SNR and ROI SNR, respectively.

### 2.12. Phase Synchronization

To estimate phase synchronization (PS) between time series of activity in ROIs, we employed three measures: imaginary part of coherency (ImCoh; Nolte et al. (2004)), lagged coherence (LagCoh; Pascual-Marqui et al. (2011)), and coherence (Coh; the absolute value of coherency). All these measures are derived from the complex coherency *c*(*f*), which can be estimated using the Fourier transform *x*(*f*) and *y*(*f*) of the time series of interest at frequency bin *f* :

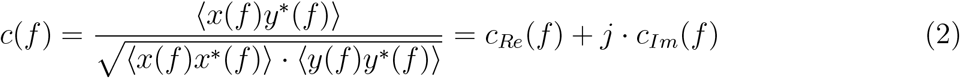

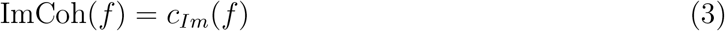

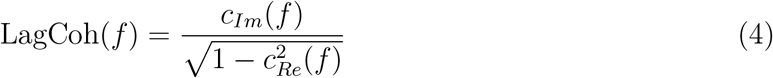

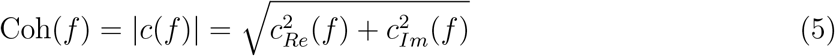

In the equations above, *j* denotes the imaginary unit, while *z*^*\**^ is a complex conjugate of *z*. Averaging over trials is denoted by < *·* >.

We considered multiple PS measures due to their complementary properties. ImCoh and LagCoh are insensitive to all zero-lag interactions, including the spurious ones caused by the volume conduction. However, it may be hard to interpret correlations between performance and PS as measured by ImCoh and LagCoh. Both measures depend on the strength and the phase lag of the interaction between neuronal populations. If ImCoh or LagCoh is correlated with performance, it is not entirely clear whether the strength or the phase lag of the interaction drives the correlation. At the same time, coherence is supposed to solely reflect the strength of an interaction, but is prone to the effects of volume conduction and might be spurious. To combine interpretability and robustness to spurious zero-lag interactions, we have considered all of these PS measures and looked at whether the observed effects are consistent between them.

We computed the PS via the Fourier transform for broadband data using the Hamming window and 1.5 s segments from different trials (frequency resolution = 0.67 Hz). First, we inspected the PS spectra in the 3-40 Hz range to confirm the presence of the peak in the frequency range of interest (9-15 Hz). For the subsequent analyses of the relationship between PS and BCI performance and changes in PS over time, we averaged the absolute values of PS measures across all frequencies in the 9-15 Hz range.

In the case of several SVD components per ROI, PS values were first computed for each pair of the SVD components, then the absolute values of PS were averaged. Furthermore, absolute PS values were averaged over within-hemisphere and across-hemisphere edges as shown in Fig. 1F, which resulted in two values (i.e., within and across-hemisphere PS) per session for each subject similar to (Vidaurre et al., 2020).

Since changes in the SNR of oscillations in the frequency band of interest lead to spurious changes in PS due to either more or less accurate phase estimation (Muthukumaraswamy and Singh, 2011), we applied a correction for SNR in the statistical analyses.

### 2.13. Multiverse Analysis

Overall, in the current multiverse analysis, we considered 24 pipelines based on all possible combinations of methods for the aforementioned processing steps (Fig. 2A). By selecting these pipelines, we aimed to assess the effects of data-independent (eLORETA, AVG-F, BB, anatomical ROIs) and task- or subject-dependent (LCMV, SVD, NB, task-based ROIs) methods on the estimated values of SNR and PS, their relationship with BCI performance, and changes over time. For each pipeline, we estimated the values of SNR as well as within- and across-hemisphere PS. Then, we tested their relationship with performance and changes over time, as described below.

### 2.14. Statistical Analysis

Statistical analysis was performed in R 4.2.2 (R Core Team, 2022). We used one-sample t-tests to analyze differences in mu power between trials corresponding to the left- and right-hand imaginary movements. Also, we used Welch’s two-sample t-test to check for group differences in performance and SNR. To assess the between-subject effects of SNR or PS on BCI performance, we correlated accuracy and a predictor variable (SNR or PS) after averaging them over all sessions for each subject. Within-subject effects of SNR and PS on accuracy as well as changes in SNR and PS over time were assessed with linear mixed-effect (LME) models using lme4 (Bates et al., 2015) and lmerTest (Kuznetsova et al., 2017) packages. The values of continuous variables were normalized before fitting the LMEs by subtracting the mean and dividing over the standard deviation. The denominator degrees of freedom in the LMEs were adjusted according to Satterthwaite’s method (Satterthwaite, 1946). P-values less than 0.05 were considered significant. The LME models that correspond to the research questions (relationship between SNR or PS and BCI performance, changes in SNR and PS over time, and effects of different processing methods on SNR and PS) are presented in Table 1. Additionally, we used linear mixed models to investigate the relationship between SNR and PS values.

**Table 1:** Linear mixed-effects models that were used for the assessment of the effects of interest. Notation X→ Y|Z corresponds to the effects of X on Y, controlled for Z. Acc., Filt., Inv., ROI, and Extr. stand for Accuracy, Filtering, Inverse Modeling, ROI Definition, and Extraction of ROI Time Series, respectively. Random slopes were added to the models as long as they converged for all of the considered pipelines. Notation (*·*)^2^ is used to show that all two-way interactions between predictors in brackets were included in the model.

| Effect | Model |  |
| --- | --- | --- |
| *Relationship between SNR and Phase Synchronization (PS) Values* |  |  |
| SNR → PS | $PS \sim 1 + SNR + (1 \vert \text{Subject})$ | (*) |
| *Relationship between SNR or PS and BCI Performance (Accuracy)* |  |  |
| SNR → Acc. | $Accuracy \sim 1 + SNR + (SNR \vert \text{Subject})$ | (*) |
| PS → Acc. | $Accuracy \sim 1 + PS + (1 \vert \text{Subject})$ | (*) |
| PS → Acc. \| SNR | $Accuracy \sim 1 + SNR + PS + (1 \vert \text{Subject})$ | (*) |
| *Longitudinal Changes in Accuracy, SNR, and PS* |  |  |
| Session → Acc. | $Accuracy \sim 1 + Session + (Session \vert \text{Subject})$ | |
| Session → SNR | $SNR \sim 1 + Session + (Session \vert \text{Subject})$ | (*) |
| Session → PS | $PS \sim 1 + Session + (1 \vert \text{Subject})$ | (*) |
| Session → PS \| SNR | $PS \sim 1 + SNR + Session + (1 \vert \text{Subject})$ | (*) |
| *Effects of the Processing Methods on the Estimated SNR and PS* |  |  |
| Methods → SNR | $SNR \sim (\text{Filt.} + \text{Inv.} + \text{ROI} + \text{Extr.})^2 + (1 \vert \text{Subject}) + (1 \vert \text{Pipeline})$ | |
| Methods → PS | $PS \sim (\text{Filt.} + \text{Inv.} + \text{ROI} + \text{Extr.})^2 + (1 \vert \text{Subject}) + (1 \vert \text{Pipeline})$ | |
| Methods → PS \| SNR | $PS \sim SNR + (\text{Filt.} + \text{Inv.} + \text{ROI} + \text{Extr.})^2 + (1 \vert \text{Subject}) + (1 \vert \text{Pipeline})$ | |
| (*) random effect of the processing pipeline (1 \| Pipeline) was added in the joint multiverse analysis. |  |  |

For the multiverse analysis, we have considered two approaches: split and joint analysis. In the split analysis, we fitted a separate mixed model for each of the pipelines and then aggregated the results in the form of a table as shown in Fig. 2D. For each pipeline, estimated correlations (between-subject effect) or regression coefficients (within-subject effect) were coded with color, and the significance of the effects was indicated by filled black dots. With this representation, one can visually inspect whether the effect is robust or specific to one of the processing methods.

In the joint analysis, we first combined the data from all pipelines and then ran the statistical analysis while including the pipeline as a random factor in the linear mixed model (see the asterisks in the rows of Table 1). This way, we obtained one result for each research question based on the combined evidence from all considered pipelines. Additionally, we calculated the consistency between pipelines as the number of pipelines that led to the same result (in terms of significance and, if significant, direction of the effect) as the joint analysis. Effects for all performance predictors and research questions were aggregated in a table as shown in Fig. 2E, where consistency is coded with color and significance is indicated by filled black dots.

Finally, we analyzed the effects of different processing methods on the estimated values of SNR and PS. Processing steps were modeled as categorical variables with Broadband, eLORETA, Anatomical ROI definitions, and 1SVD as reference levels for Filtering, Inverse Modeling, ROI definitions, and Extraction of ROI time series, respectively. To assess the impact of the combination of methods on the estimated effects of SNR and PS, we included fixed effects of all processing steps and all two-way interactions in the model.

For all research questions, we applied the Bonferroni correction for multiple comparisons (*m* = 6) since we considered two options (within- and across-hemisphere) for three PS measures (ImCoh, LagCoh, and coherence). We did not apply correction for multiple comparisons due to having 24 pipelines, since we assumed that each pipeline is equally likely to be selected for the estimation of PS. Instead, the split analysis was performed to investigate which of the individual pipelines led to a significant result.

## 3. Results

### 3.1. Performance improved over time and did not differ between groups

The average accuracy of BCI control increased from 64.3 % in the baseline session to 76.5 % in the last session (Fig. 3A). Longitudinal changes were assessed with a linear mixed model, and the effect of session was statistically significant (*β* = 0.12, *t*(57.9) = 2.8, *p* = 0.006, 95% CI: [0.03, 0.20]). As shown in the previous analyses of the same dataset (Stieger et al., 2020), there were no significant differences in the mean accuracy between MBSR (70.99 %) and control (71.08 %) groups: *t*(57.5) = − 0.02, *p* = 0.98, Cohen’s *d* = − 0.006, 95% CI of the difference [−0.07, 0.07] (Fig. 3B). The mean accuracy of all participants was 71.03%. At the same time, the intra-individual variability of performance was quite considerable (Fig. 3C). We used linear mixed models to account for this variability in the current analysis.

**Figure 3:**
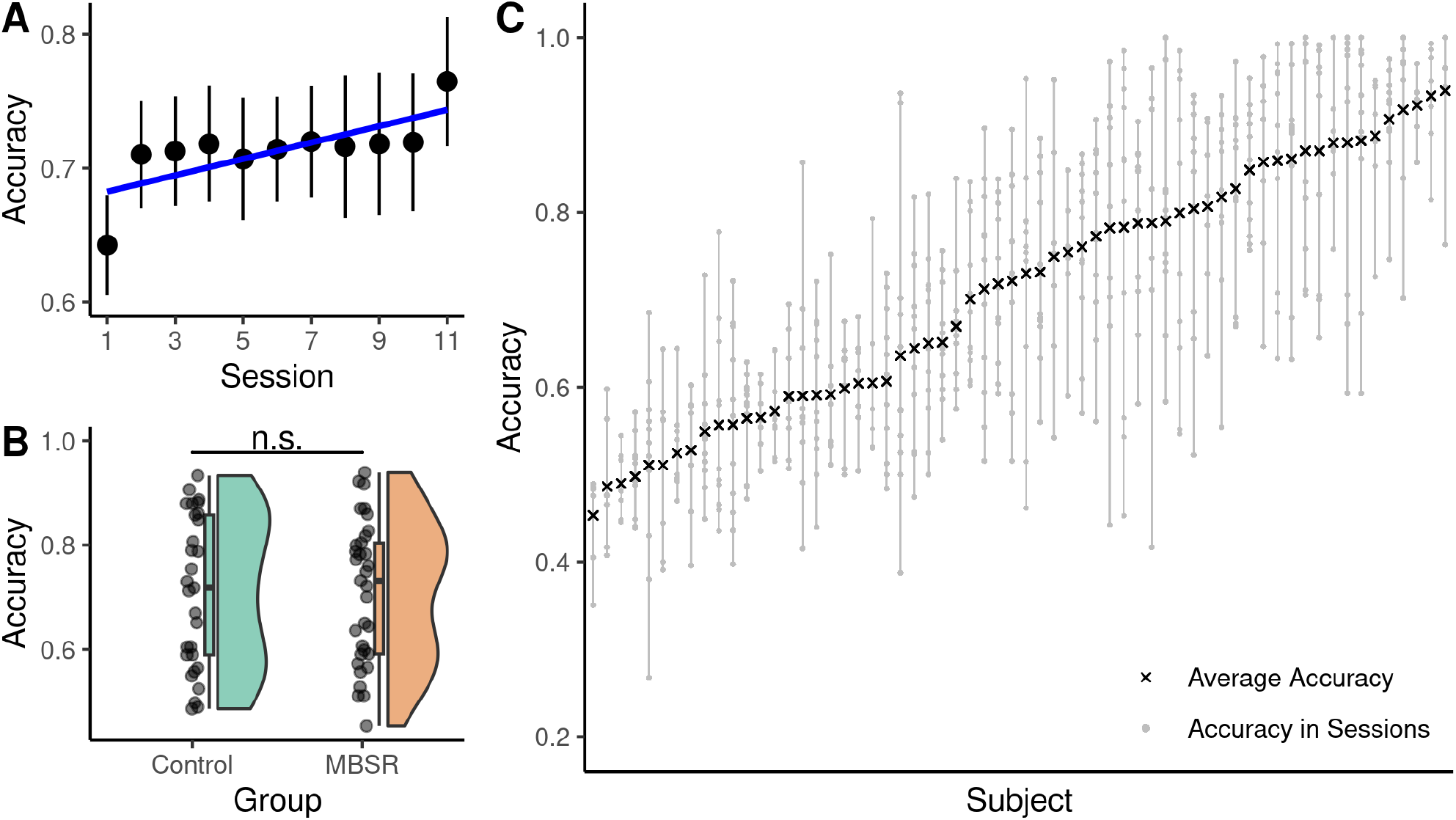
Performance of the participants improved throughout the BCI training and showed high between- and within-subject variability. (A) Dynamics of group-average performance reflect improvement over the course of the training. Error bars reflect the standard error of the mean. (B) No difference in average performance in the horizontal cursor control task was observed between the MBSR (mindfulness-based stress reduction) and control groups. (C) High variability of performance in the individual sessions was observed and accounted for in the analyses. Subjects are ordered according to their average accuracy. Vertical bars depict subject-specific ranges of accuracy.

### 3.2. Sensorimotor ROIs contained the majority of task-relevant sources

For some of the source space analysis pipelines, we identified the task-relevant sources by fitting CSP to distinguish between imaginary movements of two hands. For this purpose, we used EEG during the target presentation interval as it showed a difference in mu power between the imaginary movements primarily over the sensorimotor areas (Fig. 1C). The resulting CSP patterns and the corresponding power spectra for left- and right-hand movements are shown in Fig. 4A and Fig. 4B, respectively. These patterns were source reconstructed with eLORETA to assess the contribution of individual sources to CSP components (Fig. 4C). Sources that exceeded the 97.5th percentile of activity strength were considered task-relevant, and Tab. S4 shows that the sensorimotor ROIs contained the highest number of selected sources. Task-relevant sources formed the resulting source mask (Fig. 4D), which was applied to the anatomical definitions of sensorimotor ROIs to obtain a task-based reduced representation (Fig. 2B).

**Figure 4:**
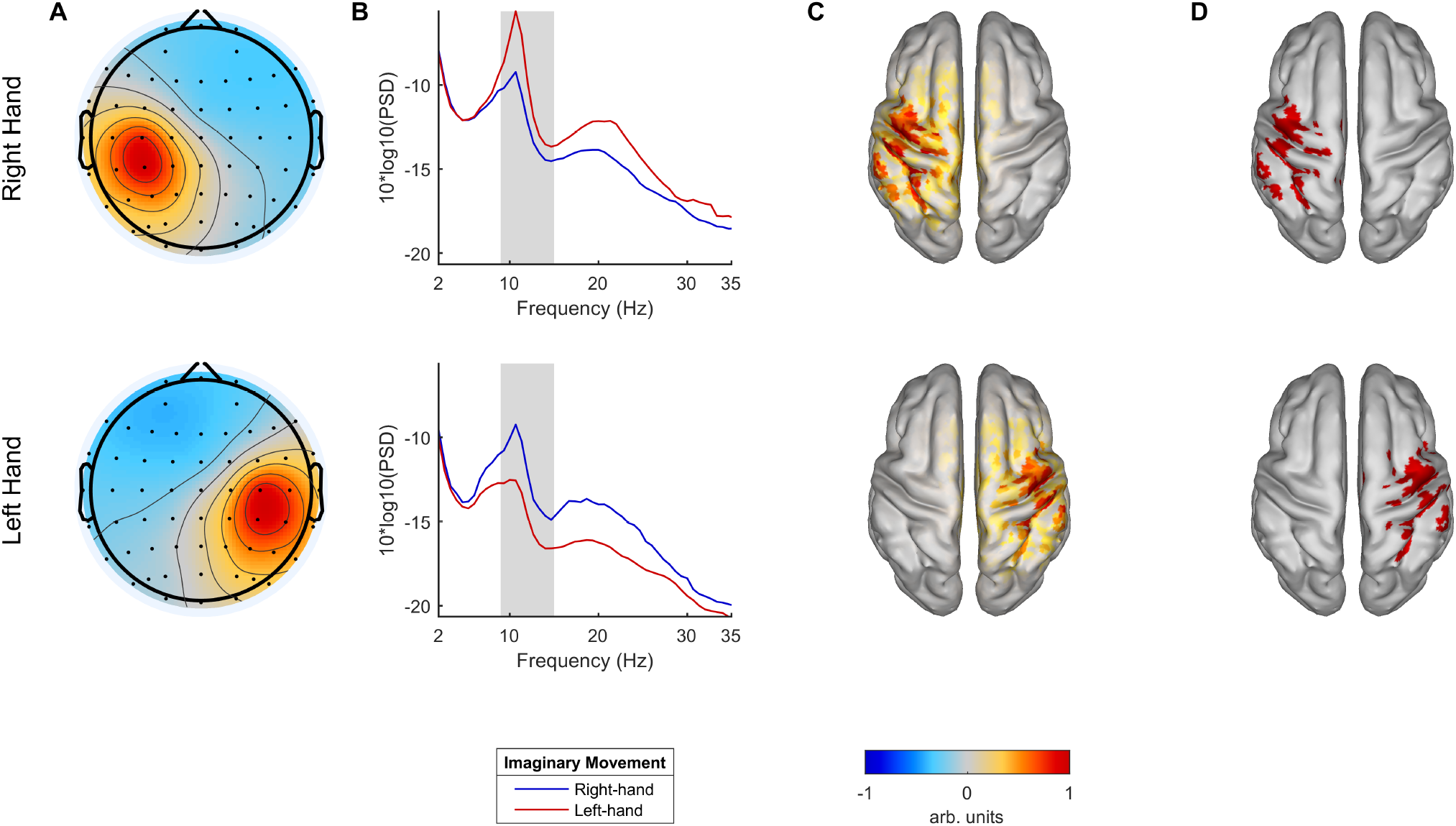
The task-relevant sources were identified through applying CSP to the EEG data during the target presentation interval after filtering in the 9-15 Hz frequency band. (A) Spatial patterns corresponding to the CSP filters that best discriminate imaginary movements of the right (upper row) and left (lower row) hands. Values were scaled to the [-1, 1] range. (B) Grand average power spectra of the CSP components corresponding to the spatial patterns from (A). The shaded area depicts the 9-15 Hz frequency band that was used to fit CSP. (C) Source reconstruction (absolute values, scaled to [-1, 1] range) of the spatial patterns from (A) with eLORETA. (D) Sources that exceeded the 97.5th percentile of activity strength were considered the most relevant for the execution of the motor imagery task.

### 3.3. Laplace SNR was correlated with BCI performance but did not change across experimental sessions

In the Laplacian-based analysis, we estimated the effects of SNR and phase synchronization (PS; see Supplementary Material, Section C) on BCI performance and their longitudinal changes. We used FOOOF to estimate average values of SNR at the Laplace-filtered channels C3 and C4. Examples of average power spectra for three representative subjects with different levels of Laplace SNR are shown in Fig. 5A. Similar to performance, Laplace SNR did not differ significantly between the participant groups as shown in Fig. 5B (*t*(59.7) = 1.09, *p* = 0.28, Cohen’s *d* = 0.28, 95% CI of the difference: [−0.63, 2.1]). Dynamics of the group-average Laplace SNR across experimental sessions are shown in Fig. 5C.

**Figure 5:**
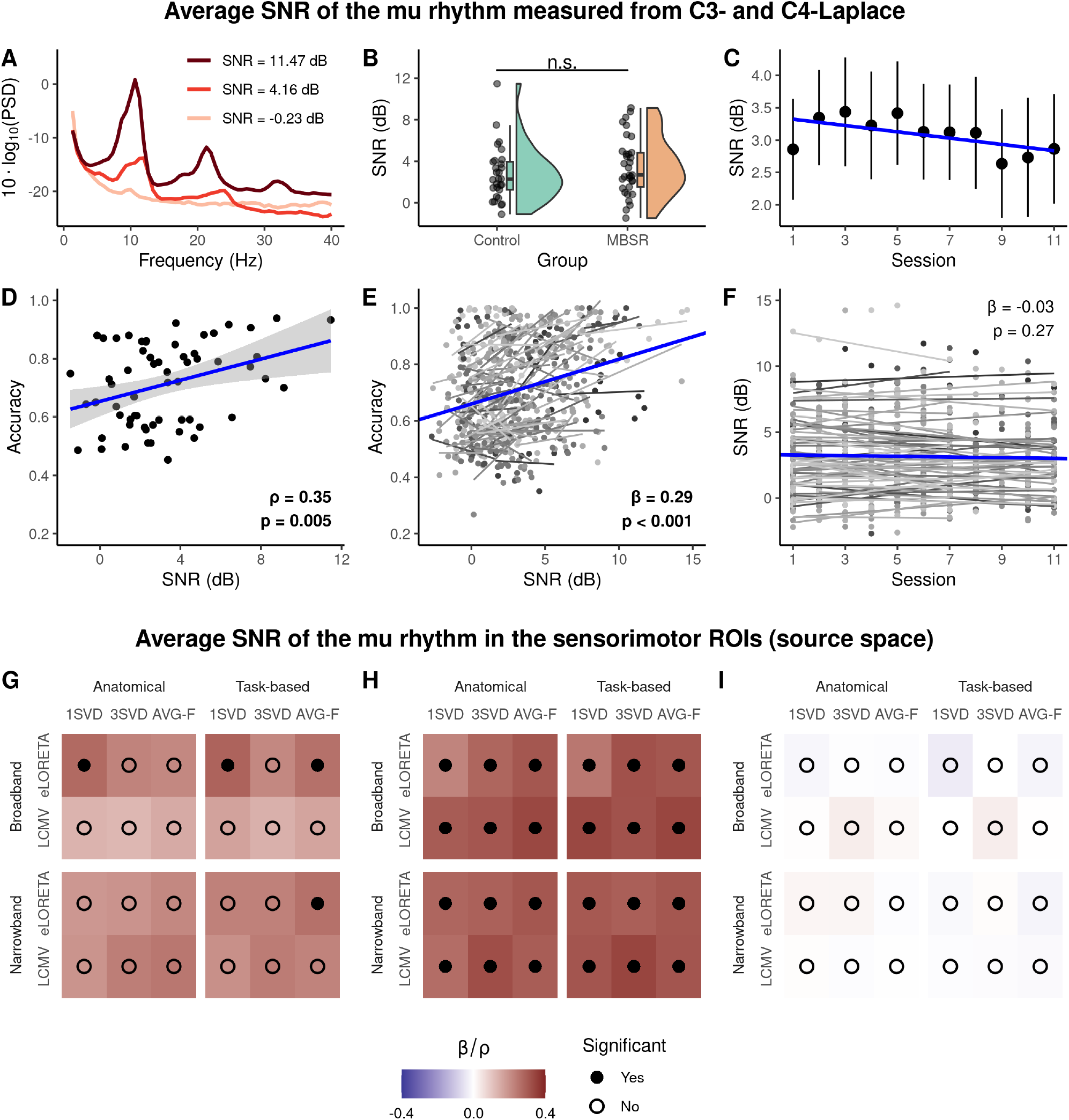
Laplace and ROI SNR showed both between- and within-subject effects on BCI performance and did not change systematically throughout the training. (A) Examples of resting-state power spectra (average of C3- and C4-Laplace over all sessions) for representative subjects with different levels of Laplace SNR. (B) The difference in SNR between groups was not significant. (C) Dynamics of group-average SNR across sessions. (D) Accuracy positively correlated with SNR after averaging over all sessions. Each point corresponds to a single participant. (E) Within-subject variability of BCI performance was related to session-to-session changes in SNR. Each point corresponds to a single session. Within-subject (gray) and group-level (blue) linear trends are shown. (F) No longitudinal changes were observed for SNR. Within-subject (gray) and group-level (blue) linear trends are shown. Note the difference in y-axis scale compared to Panel C. (G) Multiverse analysis of between-subject correlation (*ρ*, coded with color) between ROI SNR and BCI performance. (H) Within-subject effect (*β*, coded with color) of ROI SNR on BCI performance in a multiverse analysis. (I) No evidence for longitudinal changes in ROI SNR was observed for all pipelines in the multiverse analysis. Fixed effect of session on ROI SNR (*β*) is coded with color.

Similar to (Blankertz et al., 2010a), we checked whether Laplace SNR was related to successful performance in the BCI training. Subject-average values of Laplace SNR were positively correlated with accuracy (*r* = 0.35, *t*(60) = 2.9, *p* = 0.005, 95% CI: [0.11, 0.55]), showing a between-subject effect of Laplace SNR on performance (Fig. 5D). Additionally, the within-subject effect of Laplace SNR on accuracy was significant, as assessed with a linear mixed model (*β* = 0.29, *t*(57.2) = 5.22, *p* < 0.001, 95% CI: [0.18, 0.40]). Figure 5E illustrates the observed within-subject effect.

Then, we investigated whether Laplace SNR changed over time due to the training, but longitudinal changes were not significant (*β* = − 0.03, *t*(51.2) = − 1.11, *p* = 0.27, 95% CI:[−0.07, 0.02]). Individual and group-level trends are shown in Fig. 5F.

### 3.4. Effects of source space SNR, but not phase synchronization, were stable in the multiverse analysis

In the source space analysis, we estimated values of ROI SNR and phase synchronization (PS) in sensorimotor brain areas and investigated their relationship to the BCI performance as well as changes throughout the training. We performed a multiverse analysis to investigate the robustness of the observed effects to the selection of the pipeline. Figures 5G and 5H show that the estimated effects of ROI SNR on accuracy were positive for all 24 pipelines both on the between- and within-subject levels, respectively. Additionally, on the within-subject level, all effects were significant. Figure 5I shows that no significant longitudinal changes in ROI SNR were observed for all considered pipelines. Overall, the results of the multiverse analysis for ROI SNR corresponded to the results for Laplace SNR and showed that the selection of the pipelines did not affect the observed effect of ROI SNR on performance and changes in ROI SNR.

For the PS, we first checked whether the grand-average spectra of within- and across-hemisphere values of PS measures show a pronounced peak in the mu frequency range. Such a peak indicates that the interaction is specific to the ongoing mu oscillations. As shown in Fig. 6A for anatomical and Fig. S6 for task-based definitions of ROIs, the peak was pronounced in most cases. However, within-hemisphere coherence estimated using the first SVD component showed almost identical values in the whole frequency range. In this case, it might occur due to the volume conduction, which equally affects all the frequencies.

**Figure 6:**
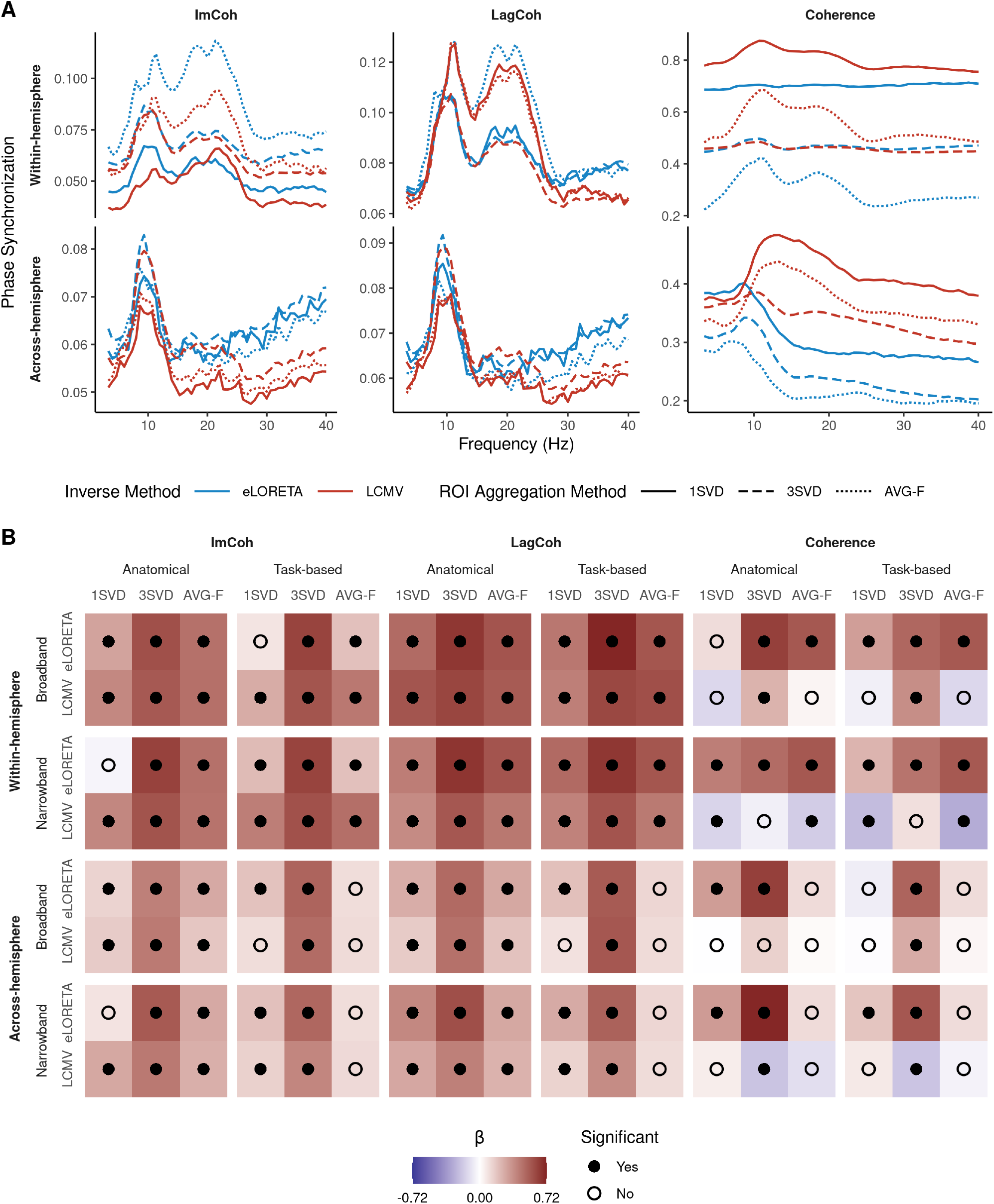
Multiverse analysis of the relationship between SNR and phase synchronization (PS) measures. (A) Grand average spectra of within- (top row) and across-hemisphere (bottom row) values of ImCoh, LagCoh, and coherence (columns: left to right) for the broadband pipelines with anatomical definitions of ROIs and different ROI aggregation methods. (B) ROI SNR showed consistent positive effects on ImCoh and LagCoh but not on coherence, both for within- and across-hemisphere PS. Fixed effect of ROI SNR on PS (*β*) is coded with color.

In line with the previous studies (Bayraktaroglu et al., 2013; Vidaurre et al., 2020), we observed a robust positive effect of ROI SNR on ImCoh and LagCoh, which are not sensitive to both spurious (caused by volume conduction) as well as genuine zero-lag interactions (Fig. 6B). In contrast, the effects of ROI SNR on coherence were less consistent between pipelines and differed in sign depending on the selection of the processing methods. Overall, these results confirm that it is necessary to account for SNR in the analyses of effects related to PS.

Then, we investigated the relationship between PS and accuracy as well as changes in PS over time. For both research questions, effects were not significant for the majority of the pipelines and PS measures (Fig. 7). Nevertheless, the pipelines that led to significant results often corresponded to a choice of a particular method at different processing steps. For example, the effects of within-hemisphere ImCoh and LagCoh on accuracy were more likely to be significant when inverse modeling was performed with an LCMV beamformer (Fig. 7A, rows 2 and 4 from the top). In this case, pipeline-specific results showed up as stripes in the visualization. A different tendency was observed for the between-subject effect of PS on accuracy (Fig. S7) as well as longitudinal changes in PS (Fig. S8): When assessing PS using coherence, significant effects were more likely to emerge than for other PS measures. Overall, the effects of connectivity on performance were not significant for the majority of the pipelines, and the direction of the effects was not consistent between different pipelines and PS measures.

**Figure 7:**
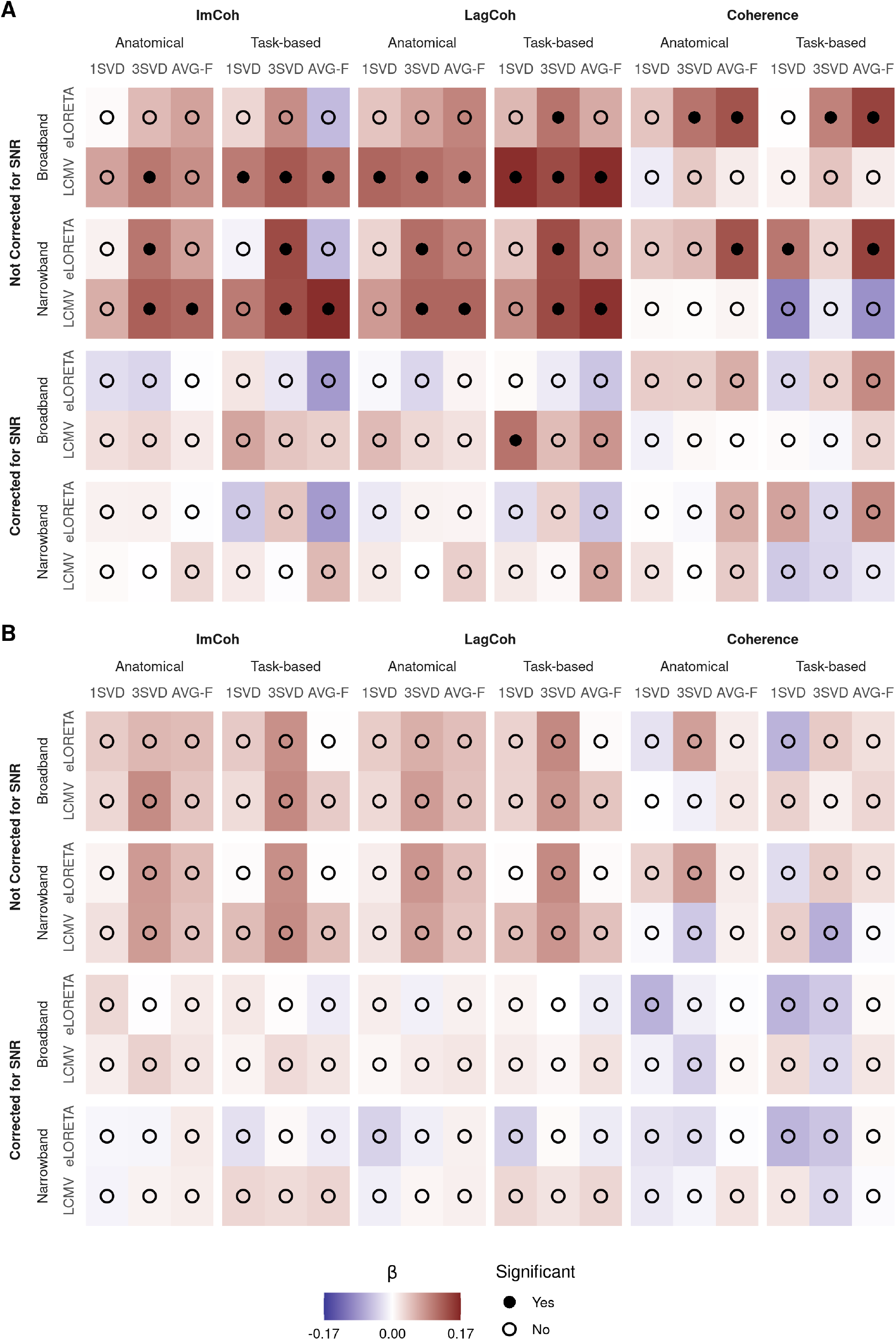
Within-subject effects of PS on BCI performance (*β*, coded with color) in the split multiverse analysis. Bonferroni correction for multiple (*m* = 6) comparisons was applied. Panels (A) and (B) correspond to within- and across-hemisphere PS, respectively.

Finally, we ran a joint analysis for all research questions by pooling together the data from all of the pipelines and fitting one linear mixed model per question (Fig. 8). Once again, the aforementioned effects of ROI SNR on accuracy and PS were significant and robust to the selection of the pipeline. Effects of ImCoh and LagCoh on accuracy were significant before correction for ROI SNR but less consistent between considered pipelines. For ImCoh, the effect on accuracy remained significant in the joint analysis after correction for ROI SNR, but none of the pipelines showed the same effect in the split analysis (when a separate mixed model is fitted for each of the pipelines). Based on the evidence from all of the pipelines, across-hemisphere LagCoh and coherence significantly increased over the course of the training. However, for LagCoh longitudinal changes were significant only in two pipelines. Statistical results are presented in Table S5.

**Figure 8:**
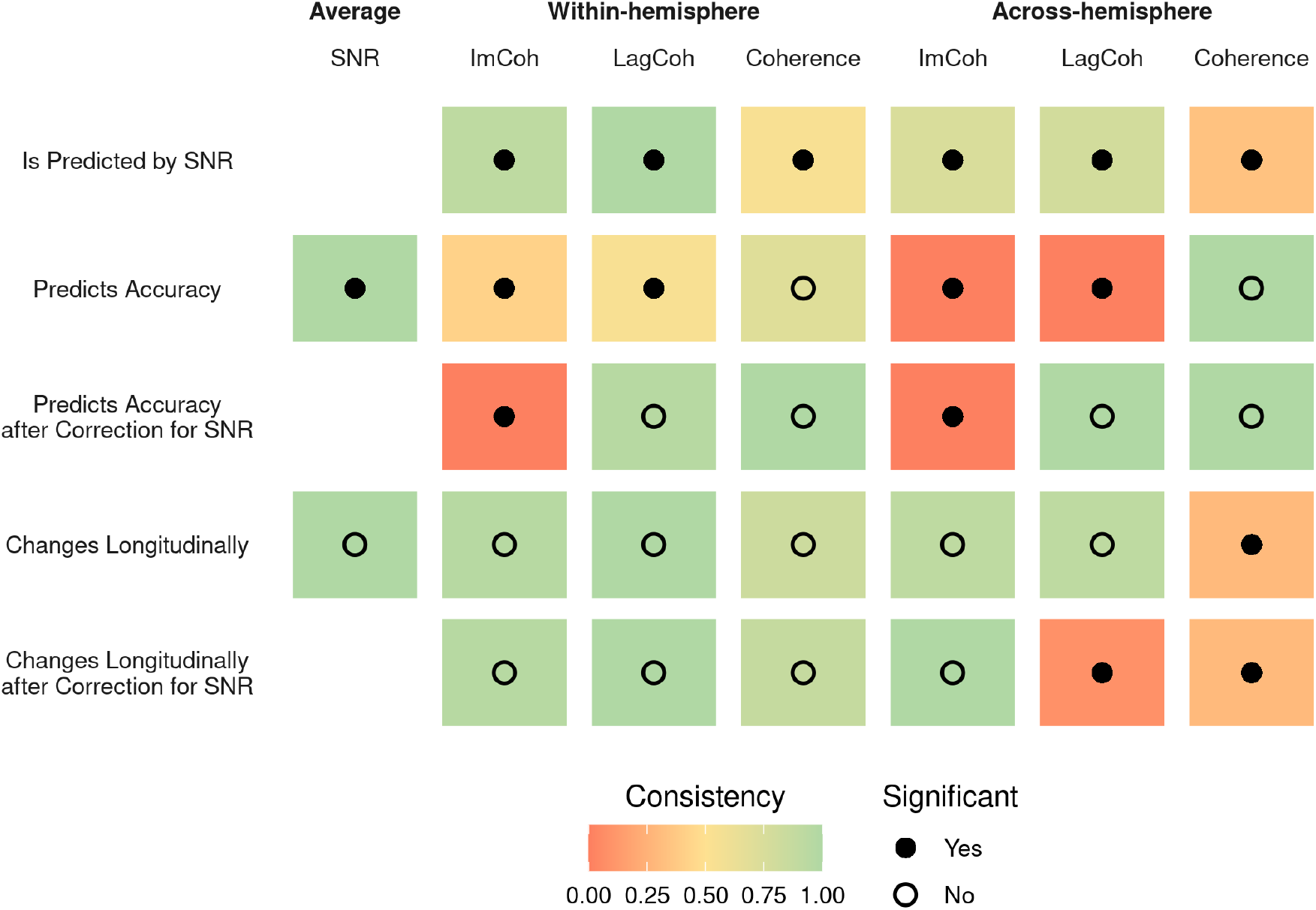
Overview of the observed within-subject effects in the joint multiverse analysis. Bonferroni correction for multiple (*m* = 6) comparisons was applied to account for several PS measures. Color codes the number of pipelines that led to the same statistical result (and, if significant, the same direction of the effect) as the joint analysis.

### 3.5. The selection of processing methods for the source space analysis affected the estimated values of SNR and PS

Effects of different methods on the estimated values of ROI SNR and PS were assessed with a linear mixed model. We included fixed effects of all processing steps and all two-way interactions in the model to investigate whether the selection of the pipeline systematically affected the estimated values of SNR and PS. Table 2 contains the estimated t-values for fixed effects of all predictors, while Table S6 lists all two-way interactions that were significant. In both tables, significant effects are highlighted in bold, and stars indicate that the effects remained significant after the correction for multiple comparisons.

**Table 2:** Summary of the observed fixed effects (t-values) of different processing methods on the estimated values of SNR and phase synchronization (PS). Significant effects are highlighted in bold, and stars indicate that the effects remained significant after Bonferroni correction for multiple (*m* = 6) comparisons. Columns correspond to different processing steps, and a positive t-value for *Y*|*X* denotes that SNR or PS was higher when *Y* was used compared to *X*. X | SNR denotes that a correction for SNR was applied. WH and AH stand for within- and across-hemisphere, respectively.

| Value | Band<br>NB \| BB | Inverse Method<br>LCMV \| eLORETA | ROI Def.<br>Task \| Anat. | ROI Method<br>3SVD \| 1SVD | ROI Method<br>AVG-F \| 1SVD |
| --- | --- | --- | --- | --- | --- |
| SNR | **3.38\*** | **6\*** | 1.22 | -0.24 | 1.29 |
| WH ImCoh | 0.01 | -1.29 | **-3.31** | **2.79** | **5.05\*** |
| WH LagCoh | 1.14 | 0.87 | -2.18 | -1.58 | 2.21 |
| WH Coherence | 0.03 | 1.53 | 2.15 | **-3.63\*** | **-4.2\*** |
| AH ImCoh | -0.13 | -0.78 | 0.41 | **3.3** | 0.86 |
| AH LagCoh | 2.03 | -0.23 | **-2.79** | **3.19** | -1.62 |
| AH Coherence | 1.57 | **2.87** | **-2.72** | -2.21 | **-2.51** |
| WH ImCoh \| SNR | -0.38 | -1.93 | **-3.31** | **2.69** | **4.69\*** |
| WH LagCoh \| SNR | -1.1 | **-3.52\*** | **-3.96\*** | -1.98 | 2.02 |
| WH Coherence \| SNR | -0.18 | 1.2 | 2.14 | **-3.73\*** | **-4.41\*** |
| AH ImCoh \| SNR | -0.74 | -1.82 | 0.14 | **3.02** | 0.54 |
| AH LagCoh \| SNR | 0.27 | **-3.1** | **-3.22** | **3.12** | -2.15 |
| AH Coherence \| SNR | 1.35 | **2.49** | **-2.91** | -2.26 | **-2.69** |

First, we observed that the values of SNR were affected by filtering and the choice of the method for inverse modeling, with the interaction of these processing steps also being significant (Fig. 9A). We investigated this result in more detail since the quality of the source reconstruction with LCMV depends on the SNR (Van Veen et al., 1997), and SNR played an important role in the previous analyses. For this purpose, we compared values of SNR within pairs of pipelines, which differed only in the method for inverse modeling. As shown in Fig. S9, LCMV led to higher SNR than eLORETA for broadband pipelines, and the difference was especially pronounced for low values of SNR. For narrowband pipelines, SNR was on average higher when eLORETA was used.

**Figure 9:**
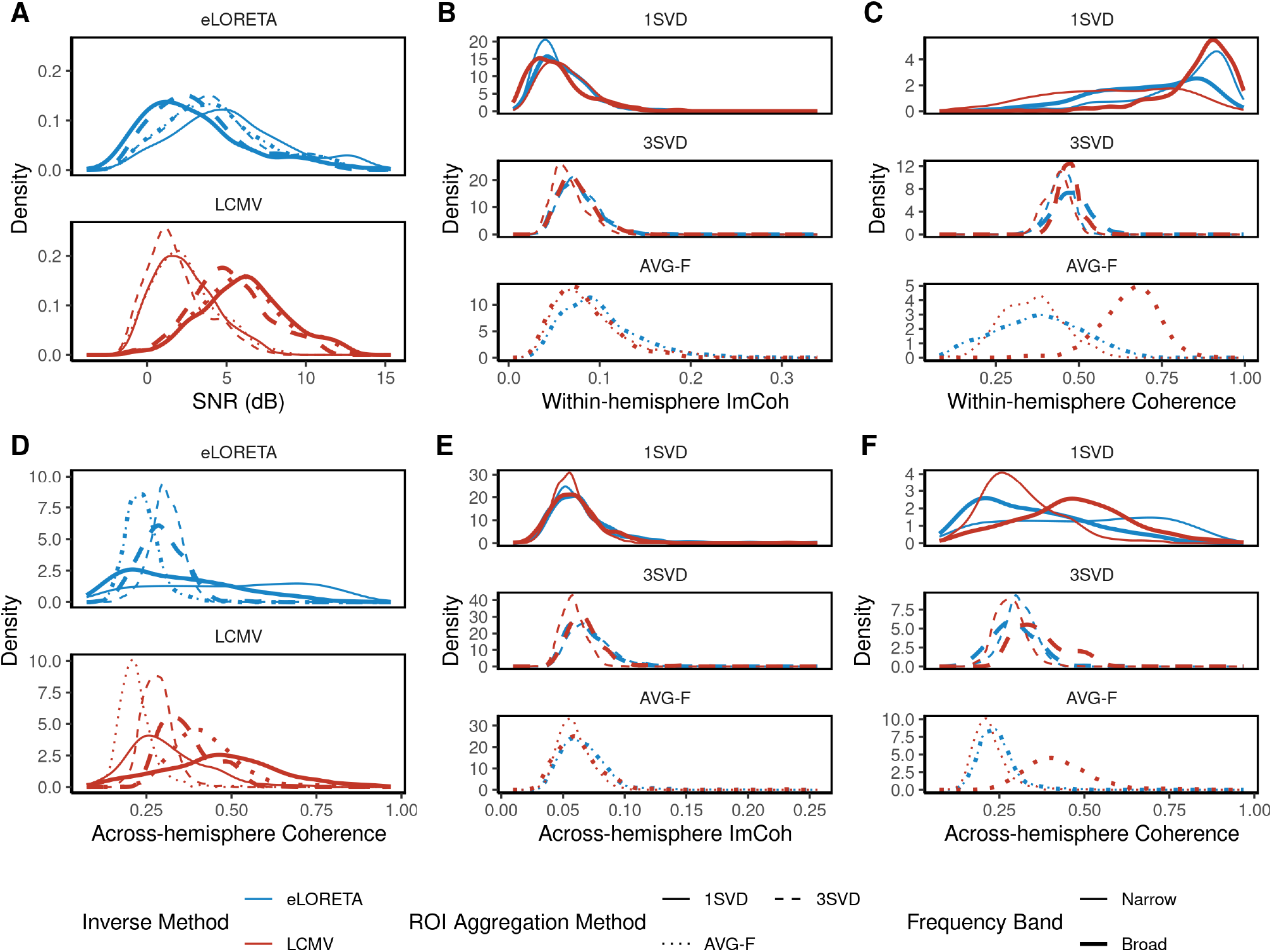
Selection of the processing methods affected estimated values of ROI SNR and PS as indicated by shifts in the empirical probability density functions. Only pipelines with anatomical definitions of ROIs are displayed. (A) SNR was affected by filtering and the choice of method for inverse modeling. (B) Within-hemisphere ImCoh was higher for 3SVD and AVG-F compared to 1SVD. (C) Method for extraction of ROI time series affected values of within-hemisphere coherence. (D) LCMV led to higher values of across-hemisphere coherence compared to eLORETA. (E) Same as B, but for across-hemisphere ImCoh. (F) Same as C, but for across-hemisphere coherence.

PS measures were affected by the selection of methods for all processing steps except filtering. When 3SVD or AVG-F were used for the extraction of ROI time series as compared to 1SVD, coherence decreased (Fig. 9, Panels C and F), while ImCoh increased compared to 1SVD (Fig. 9, Panels B and E). Task-based definitions of ROIs led to a decrease in within-hemisphere ImCoh and LagCoh as well as across-hemisphere LagCoh and coherence. Additionally, for within-hemisphere PS, there was a significant interaction between the ROI definition and the method for extraction of ROI time series. In particular, pipelines with 3SVD were less affected by the definition of the ROI (Fig. S10). Finally, LCMV led to smaller values of LagCoh and higher across-hemisphere coherence than eLORETA (Fig. 9D).

## 4. Discussion

In the current study, we investigated the role of SNR and phase synchronization (PS) of the mu rhythm in sensorimotor brain areas as predictors of BCI performance in a multi-session training using a publicly available dataset (Stieger et al., 2021). The dataset contained EEG recordings from a multi-session BCI training based on a cursor control paradigm. The performance of the participants was assessed with the accuracy of completed trials and improved significantly for some but not all participants throughout the training. Overall, the mean accuracy was comparable to other BCI studies and similar to the 70% threshold, which is commonly used to identify good performers (e.g., in Sannelli et al. (2019) and Leeuwis et al. (2021)). While the increase in group-average performance was not prominent between sessions 2 and 10, a considerable level of intra-individual variability of performance was observed. We used linear mixed models to account for this variability and investigate the relationship between SNR, PS, and BCI performance, as well as longitudinal changes in SNR and PS due to training. We performed the analysis in sensor space using the surface Laplacian and in source space by combining several processing pipelines in a multiverse analysis. In the following, we discuss the results of the study and their prospective applications.

### 4.1. SNR in the context of sensorimotor BCI training

Previous studies have shown that the signal-to-noise ratio of the mu rhythm estimated at the Laplace-filtered channels C3 and C4 correlated with the accuracy of BCI control (Blankertz et al., 2010a; Acqualagna et al., 2016; Sannelli et al., 2019). We also observed a positive correlation between Laplace SNR and accuracy after averaging over all sessions, which reflects a between-subject effect of SNR on performance. Additionally, we observed a within-subject effect of Laplace SNR on accuracy. That is, not only do participants with a higher SNR of the mu rhythm tend to perform better, but the same participant tends to perform better on the days when SNR is higher as well.

In general, larger SNR is associated with stronger lateralization of the mu rhythm during imaginary movements, leading to a higher classification or control accuracy (Maeder et al., 2012). Our results show that this finding, previously observed primarily for the single experimental sessions, generalizes to longitudinal settings and has two important consequences. First, changes in overall performance should be controlled for the changes in SNR to make conclusions about other possible neurophysiological factors. Second, experimental adjustments leading to an increase in SNR might also translate to a performance improvement.

It is important to note that SNR might affect BCI performance at least in two different ways. On the one hand, participants with low SNR of the mu rhythm might not be able to perform vivid imaginary movements and modulate their brain activity strongly enough. In this case, “quasi-movements” (i.e., real movements minimized to such an extent that they cannot be detected by objective measures) could be used to train participants to perform the motor imagery better (Nikulin et al., 2008). On the other hand, high SNR of the mu rhythm might translate into a more reliable feedback signal, which would in turn allow participants to train the imaginary movements more efficiently. If this is the case, training for participants with low SNR of the mu rhythm could be based on other features of brain activity that might provide a higher SNR. For example, Tao et al. (2021) have shown that motor imagery led to a decrease in inter-trial phase coherence during steady-state electrical stimulation of the median nerve.

Moreover, there is still a considerable amount of unexplained variance in BCI performance, which could be attributed to other psychological (such as motivation or concentration) and lifestyle (sports or musical instrument training) factors. These factors remain a subject of extensive research in the BCI community (Hammer et al., 2012; Jeunet et al., 2015) and could also be manipulated to improve BCI performance.

Furthermore, we investigated whether Laplace SNR itself could change throughout sensorimotor BCI training but observed no evidence of longitudinal changes. This result could be related to the structure of the cursor control tasks. Typically, post-effects of BCI or neurofeedback are observed when the whole training is based on a fixed direction of modulation of brain activity, for example, up-regulation of alpha power (Zoefel et al., 2011). In contrast, cursor control tasks in the analyzed dataset always contained trials with opposite directions of modulation of the mu rhythm (left- vs. right-hand imaginary movements or motor imagery vs. relaxation). Therefore, on average, task-related modulation of the mu rhythm may not have a cumulative effect across many sessions. In addition, Popov et al. (2023) have also reported an excellent (ICC = 0.83) test-retest reliability of the periodic component of the alpha power in the sensorimotor regions. While this finding goes in line with the absence of longitudinal changes in SNR in the analyzed dataset, there still was a within-subject effect of SNR on BCI performance. This result could be explained if SNR is a trait feature that is affected by measurement-related effects (e.g., different placement of the electrodes) on different training days. Nevertheless, measurement-related effects could, in turn, make the detection of longitudinal changes in SNR harder.

The absence of longitudinal changes in SNR is critical for discussing changes in other measures that were shown to be correlated with SNR such as phase synchronization or long-range temporal correlations (Samek et al., 2016; Vidaurre et al., 2020). Since a decrease in SNR typically leads to the attenuation of the aforementioned measures, their changes (e.g., due to learning, arousal, etc.) should be controlled for the concurrent changes in SNR.

In our study, both the positive effect of SNR on accuracy and the absence of longitudinal changes in SNR were robust to the selection of the processing steps in the multiverse analysis, as the results were the same for all of the considered pipelines. Taken together with all the existing evidence for the role of SNR in BCI training, this result might suggest that the effect of SNR on accuracy is strong enough to overcome the variability in the estimation of SNR across different pipelines.

### 4.2. Phase synchronization in the context of sensorimotor BCI training

In the current study, we analyzed three linear PS measures to combine the interpretability of coherence (as it reflects the strength of interaction) and robustness to zero-lag interactions provided by ImCoh and LagCoh. The estimation of PS was performed in the source space, and several processing pipelines were combined in a multiverse analysis to assess the variability of the PS values and associated statistical effects. For most pipelines, we observed a peak in the mu range of the PS spectra, which reflects an interaction that is specific to mu oscillations.

In line with several previous studies (Bayraktaroglu et al., 2013; Vidaurre et al., 2020), we observed a positive correlation between the values of SNR and phase synchronization. On the one hand, higher SNR improves phase estimation and may spuriously lead to higher values of PS (Muthukumaraswamy and Singh, 2011). On the other hand, a higher PS between two neuronal populations is likely to co-occur with a higher level of synchronization within the populations, which would be manifested in higher SNR values (Schneider et al., 2021). Most likely, both factors contribute to a positive correlation between SNR and PS values. This correlation was very robust to the selection of the pipeline for ImCoh and LagCoh, which are not sensitive to zero-lag spurious interactions due to volume conduction. Effects of SNR on coherence were less consistent, which could be related to the remaining spatial leakage (i.e., signal mixing), especially in the case of nearby regions within the same hemisphere. Overall, our findings confirm that it is necessary to account for changes in SNR when analyzing phase synchronization.

We observed a significant positive within-subject effect of within- and across-hemisphere ImCoh and LagCoh on BCI performance. It was significant in the joint analysis and for a few separate pipelines in the split analysis. While this finding goes in line with the results of (Vidaurre et al., 2020), we observed no evidence for a between-subject effect (Fig. S7), which could serve as a direct replication. After correction for SNR, the effect of ImCoh but not LagCoh on BCI performance remained significant in the joint multiverse analysis. At the same time, the same effect could not be detected by any of the considered pipelines in the split analysis. Therefore, our results suggest that phase synchronization was not related to BCI performance in the analyzed dataset. While motor imagery leads to a modulation of amplitude (ERD/ERS), it might not necessarily require phase synchronization as strongly as other tasks involving precise bilateral coordination (Shih et al., 2021). However, this result may also be caused by differences in the calculation of real-time feedback. In the dataset analyzed by Vidaurre et al. (2020), feedback is calculated using a CSP-based spatial filter, which is optimal for extracting task-related differences in power. In the analyzed dataset, surface Laplacian is applied to channels C3 and C4 for spatial filtering, and the resulting signal may still contain contributions from other sources, e.g., occipital alpha rhythm (Blankertz et al., 2010a).

Despite not showing high consistency between pipelines (Fig. S8), an increase in across-hemisphere coherence throughout the training was significant in the joint multiverse analysis. This result could speak in favor of the optimization of the interaction between motor areas due to the training. However, since the same effect was only observed in two pipelines for LagCoh and not at all for ImCoh, there is not enough evidence or statistical power to conclude that this increase is driven by a genuine neural interaction.

Overall, the findings related to phase synchronization were not as robust to the selection of the pipeline as they were for SNR. Hence, along with the recommendation from Mahjoory et al. (2017), it is necessary to include at least several analysis pipelines to account for the between-pipeline variability of PS values.

### 4.3. Effects of the processing methods on the estimated values of SNR and PS

The multiverse analysis also allowed us to compare SNR and PS values that were obtained by applying different combinations of methods for source space analysis to the same data. Since there is no ground truth available for real data, this comparison does not allow us to determine which methods work better or worse (Feuerriegel and Bode, 2022). Nevertheless, below we describe several observations that could be validated in simulations and used in future studies.

#### Inverse Modeling

As shown in Fig. S9, for broadband pipelines, SNR was higher on average when LCMV was used for inverse modeling compared to eLORETA. Since LCMV is a data-driven approach, it might better adapt to different subjects and sessions and thereby extract oscillatory activity with higher SNR than eLORETA. Surprisingly, the difference in SNR between pipelines with LCMV and eLORETA was especially prominent for low values of SNR. However, it is not clear whether the improvement in the SNR of the extracted signal is due to better extraction of activity from the investigated ROIs or the remaining spatial leakage from other ROIs. After correction for SNR, LCMV led to a decrease in LagCoh and an increase in across-hemisphere coherence compared to eLORETA (Fig. 6A). Previous studies (Mahjoory et al., 2017; Pellegrini et al., 2023) also observed the impact of the inverse method on the estimated PS values. While the reasons behind this effect are not clear, it is important to note that the selection of the inverse method also played a role in the split multiverse analysis. In particular, the effects of within-hemisphere ImCoh and LagCoh on BCI performance were significant only for pipelines that included LCMV (Fig. 7A).

#### Extraction of ROI Time Series

ROI time series were obtained by aggregation of time series of individual sources within the ROI, and the selection of the aggregation method affected most PS measures. In particular, for the within-hemisphere case, the first SVD component seemed to capture the remaining effects of volume conduction to a great extent, as indicated by the lack of a peak in the spectra of coherence (Fig. 6A) and the values of coherence that are very close to 1 (Fig. 9C). In contrast, when three SVD components were used for the calculation of the PS, a peak in the spectra was present, and coherence was generally lower, while ImCoh had higher values. This result might be caused by the averaging of pairwise connectivity values between different SVD components, which is more likely to result in a non-zero phase lag. Still, by including more than one component per ROI in the analysis, one might ensure that a genuine interaction between ROIs is captured. This observation goes in line with the recommendation to consider 3-4 SVD components per ROI from (Pellegrini et al., 2023). Averaging with sign flip also seemed to capture the remaining effects of volume conduction less, as reflected by lower coherence and higher ImCoh.

#### Filtering

Fig. S11 shows that for pipelines that included eLORETA as the method for inverse modeling, SNR was higher when the data were filtered in a narrow band. Since eLORETA is a data-independent method, it is not affected by the frequency range of the data. Instead, filtering matters for SVD being the only data-dependent method in the subsequent steps of the pipelines. This result supports the observation by Chalas et al. (2022) that SVD prioritizes low frequencies that explain more variance in the M/EEG signals. At the same time, the effects of filtering on all PS measures were not significant before and after controlling for SNR.

#### ROI Definition

We investigated whether reducing anatomical definitions of ROIs to a subset of task-relevant sources could make the estimated SNR and PS values even more task-specific. The definition of the ROI played a different role in the estimation of within-hemisphere and across-hemisphere PS. In the within-hemisphere case, the task-based definition reduced the size of the ROIs and variability in the reconstructed time series of individual sources. Thereby, the effects of volume conduction became pronounced even stronger (higher coherence and lower ImCoh). In the across-hemisphere case, the distance between task-based ROIs was higher than between the anatomical ones, and the observed decrease in coherence could reflect less pronounced volume conduction.

#### Interactions

Although we included all possible two-way interactions between processing steps in the model, only two of them were significant after correction for multiple comparisons (Tab. S6). First, there was a significant interaction between filtering and inverse modeling steps, which implies that there was a difference between PS and SNR values depending on whether eLORETA and LCMV were fit to broadband or narrowband data. In the analyzed dataset, narrowband LCMV led to lower SNR for all combinations of subsequent processing steps (Fig. S11). Second, for within-hemisphere PS values, there was a significant interaction between ROI definition and method for extraction of ROI time series. When three SVD components are used to describe activity in each ROI, the definition of the ROI (anatomical or task-based) seems to play a smaller role than for averaging with sign flip or a single SVD component (Fig. S10).

Overall, all processing steps affected SNR or some of the PS measures, highlighting the importance of considering several pipelines for source space analysis. Both data-dependent methods (LCMV and SVD) led to different results when fit to broadband and narrowband data. The definition of the ROI and the method for extraction of ROI time series seemed to make the biggest difference in terms of remaining volume conduction but also affected PS measures that are insensitive to zero-lag interactions.

### 4.4. Limitations

The current analysis was limited to four sensorimotor ROIs and did not include the whole-brain connectivity patterns, for example, as performed in (Corsi et al., 2020). This selection was based on previous studies showing that motor imagery BCI primarily leads to activation of the sensorimotor areas that we analyzed (Nierhaus et al., 2021). These ROIs contained the highest amount of task-relevant sources in the analyzed dataset as well (Tab. S4), thereby additionally validating the selection. As described before, there are several open questions regarding the estimation of PS, correlation between PS and behavior, correction for the effect of SNR and PS, and interpretation of the results. Analyzing only selected ROIs made it feasible to address these challenges by considering several options for each question.

There also exist other methods that were not included in the multiverse analysis to ensure computational feasibility, e.g., dynamic imaging of coherent sources (DICS; Gross et al. (2001)) for inverse modeling or fidelity weighting (Korhonen et al., 2014) for aggregation of ROI time series. However, the amount of pipelines considered in the current analysis already provides additional insights compared to a single pipeline. Still, it is important to keep in mind that even if similar results are obtained with multiple pipelines, it does not directly imply the genuineness of these results.

The final limitation is related to the longitudinal analysis. While the group-level improvement in performance was significant, group-average accuracy was similar across most sessions, which might reflect little evidence of training effects. Nevertheless, we utilized the observed within-subject variability and employed linear mixed models to estimate the effects of interest.

### 4.5. Future Directions

In the current study, we focused on predicting the BCI performance from the dynamics of mu rhythm in sensorimotor areas during the inter-trial interval, thereby testing the existing resting-state findings (Blankertz et al., 2010a; Vidaurre et al., 2020) in a longitudinal setting. Future studies could consider using task data and investigating other brain regions — for example, connectivity between visual and sensorimotor brain areas could reflect the processing of the visual feedback during motor imagery. Moreover, combining datasets with several BCI paradigms (Chevallier et al., 2024) could provide additional insights into feedback processing that generalize beyond a single paradigm (in this case, motor imagery).

Multiverse analysis seems to be a promising approach for incorporating the variability between different processing pipelines into the analysis. In the context of M/EEG connectivity, it could be interesting to evaluate the robustness of amplitude coupling measures (Hipp et al., 2012; Brookes et al., 2012) and network measures of brain connectivity (Rubinov and Sporns, 2010) to the selection of the pipeline for source space analysis.

## 5. Conclusion

Overall, we observed that SNR affected BCI performance both on the between- and within-subject levels: Participants with higher SNR tended to perform better, and the same participant also tended to perform better on the days when SNR was higher. Therefore, interventions that are suitable for increasing SNR might lead to an improvement in performance. Additionally, multiverse analyses allowed us to analyze the robustness of the investigated effects to the selection of the pipeline for source space analysis. The results suggest that SNR was a primary factor of the observed performance variability as it robustly predicted accuracy and covaried with phase synchronization (PS). On the contrary, the effects of PS became non-significant after controlling for SNR and were less consistent across different pipelines. We observed no evidence of longitudinal changes in SNR and only weak evidence of an increase in the across-hemisphere coherence during the training. At the same time, SNR and PS values were significantly affected by the selection of the pipeline for source space analysis. Therefore, it is necessary to include several pipelines in the analysis to assess how robust the observed effects are and how high the between-pipeline variability is. This paper can serve as a template for future multiverse analyses as it represents an end-to-end fully repeatable pipeline from raw data to the publishable report, and all the underlying data and scripts are publicly available.

## Data and Code Availability

EEG recordings are publicly available at https://doi.org/10.6084/m9.figshare.13123148, and a detailed description of the dataset is provided in (Stieger et al., 2020, 2021). Analysis scripts are available at https://github.com/ctrltz/bci-brain-connectivity. Preprocessing data that are necessary to reproduce the analysis (indices of bad trials, channels, and ICA components; ICA weights, excluded sessions, etc.) are available at https://osf.io/tcvyd.

## CRediT Authorship Contribution Statement

**Nikolai Kapralov:** Conceptualization, Data curation, Formal analysis, Investigation, Methodology, Software, Visualization, Writing – original draft, Writing – review & editing. **Mina Jamshidi Idaji:** Validation (Code Review), Writing – review & editing. **Tilman Stephani:** Methodology, Validation (Code Review), Writing – review & editing. **Alina Studenova:** Validation (Code Review), Writing – review & editing. **Carmen Vidaurre:** Software, Writing – review & editing. **Tomas Ros:** Writing – review & editing. **Arno Villringer:** Funding acquisition, Supervision, Writing – review & editing. **Vadim Nikulin:** Conceptualization, Investigation, Methodology, Project administration, Supervision, Writing – Original Draft, Writing – review & editing.

## Acknowledgments

We would like to thank the authors of the dataset for making it publicly available and James Stieger in particular for providing additional information about the dataset on request. C.V. was funded by the Spanish Ministry of Science, Innovation and Universities, ref. number PID2020-118829RB-I00.

