## Supplementary Material for "Sensorimotor brain-computer interface performance depends on signal-to-noise ratio but not connectivity of the mu rhythm in a multiverse analysis of longitudinal data"

[1] Now at the Donders Institute for Brain, Cognition and Behaviour, Radboud University, Nijmegen, The Netherlands.

### A. Comparison of online accuracy in different cursor control tasks

We compared the performance of participants in three cursor control tasks using Pearson's correlation coefficient (Fig. S1). Both between-subject and median within-subject correlations were high (Tab. S1), suggesting that participants performed similarly in all tasks. Still, differences in the underlying mechanisms might be observed between tasks as shown by Stieger et al. (2020). In particular, successful performance in the second task (cursor control based on total alpha power) was associated with widespread changes in alpha power during relaxation, while the first task (cursor control based on alpha lateralization) was more tightly related to the sensorimotor areas.

| Tasks | Between-subject | | Within-subject |
|---|---|---|---|
| | Correlation | 95 % CI | Median correlation |
| Task 1 ∼ Task 2 | 0.63 | [0.46, 0.76] | 0.44 |
| Task 1 ∼ Task 3 | 0.86 | [0.78, 0.92] | 0.53 |
| Task 2 ∼ Task 3 | 0.78 | [0.66, 0.86] | 0.65 |

Table S1: Between-subject and within-subject correlations of performance in the cursor control tasks.

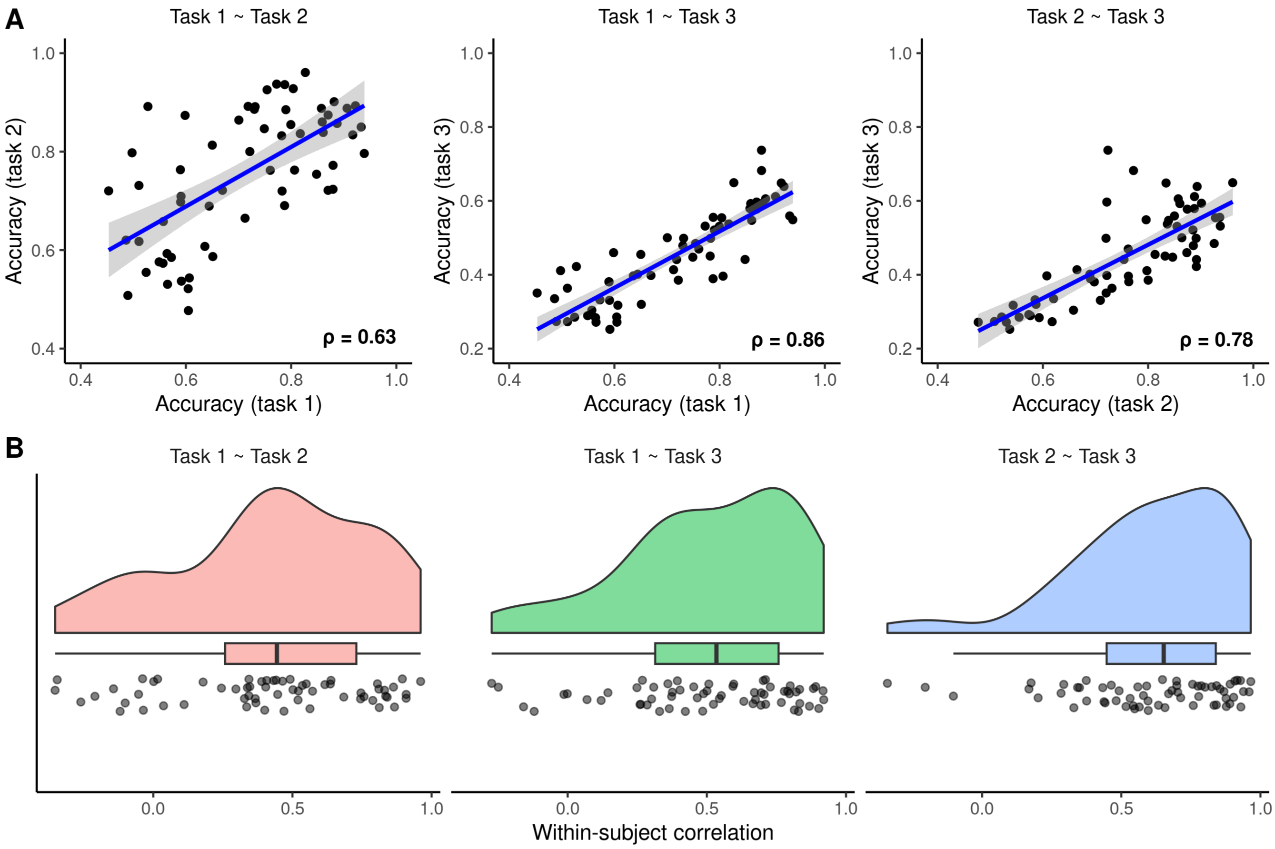


Figure S1: Comparison of performance in the cursor control tasks. (A) Between-subject comparison. Each point corresponds to a single subject and reflects average accuracy over all sessions. (B) Within-subject comparison. Each point corresponds to a single subject and reflects correlation of this subject's performance in different sessions for each pair of tasks.

### B. Comparison of online accuracy and offline estimates of performance

#### *B.1. Methods*

We used a combination of CSP (Common Spatial Pattern; Ramoser et al. (2000)) and regularized LDA (Latent Discriminant Analysis; Friedman (1989)) to distinguish between trials corresponding to left- and right-hand imaginary movements. The classification was performed using EEG data from three time windows (Fig. 1B, main text; rest: [0.49, 1.99] s of the inter-trial interval, target: [0.49, 1.99] s of the target presentation interval, feedback: [-1.51, -0.01] s relative to the end of the feedback interval) after filtering in 9-15 Hz frequency band. For both CSP and rLDA, we used implementations from the BBCI toolbox (Blankertz et al., 2016).

For each time window, we first fitted CSP and selected three components for each type of the imaginary movement (i.e., three components with the smallest and three with the largest eigenvalues). For each CSP component, we estimated the variance of the corresponding time series in the time window of interest. Then, we trained a regularized LDA using the log-variance values as features. The regularization parameter of LDA was set to 0.05. We estimated *offline* accuracy and area under the receiver operating characteristic curve (AUC) of the classifier for each subject and training session separately using 6-fold cross-validation. To compare online accuracy and offline estimates of performance, we used Pearson's correlation coefficient.

#### *B.2. Results*

The results of the analysis are presented in Fig. S2. During the inter-trial interval (labeled as rest), both accuracy (Fig. S2-A) and AUC (Fig. S2-D) were not significantly different from chance level of 0.5 after correction for multiple comparisons. At the same time, both accuracy and AUC increased and became significantly higher than chance level when the data during the target presentation and feedback intervals were used for training the classifier. The detailed statistical results are presented in the Table S2.

For comparison with the online accuracy, we selected offline estimates based on EEG data from the feedback interval. We observed high correlation between online accuracy and offline performance metrics (accuracy: $\rho = 0.71, 95\%$ CI: $[0.56, 0.81]$, AUC: $\rho = 0.7, 95\%$ CI: $[0.55, 0.81]$) on the between-subject level (Fig. S2-B and -E). On the within-subject level (Fig. S2-C and -F), the correlation was subject-specific but still relatively high for the majority of participants (median within-subject correlations for accuracy and AUC were 0.57 and 0.55, respectively). Overall, these results suggest that the main findings of the paper should still hold if offline accuracy or AUC is used as the metric of performance.

| Metric | Period | Mean | t-value | df | p-value | 95% CI |
|---|---|---|---|---|---|---|
| Accuracy | rest | 0.51 | 2.04 | 61 | **0.046** | $[0.50, 0.51]$ |
| | target | 0.66 | 11.78 | 61 | **<0.001*** | $[0.63, 0.68]$ |
| | feedback | 0.69 | 12.42 | 61 | **<0.001*** | $[0.66, 0.72]$ |
| AUC | rest | 0.51 | 2.19 | 61 | **0.032** | $[0.50, 0.52]$ |
| | target | 0.70 | 12.71 | 61 | **<0.001*** | $[0.67, 0.73]$ |
| | feedback | 0.73 | 13.57 | 61 | **<0.001*** | $[0.70, 0.77]$ |

Table S2: Summary of the observed differences between CSP-based ('Mean' column) and chance level (0.5) values of accuracy and AUC in three time windows of interest (rest: [0.49, 1.99] s of the inter-trial interval, target: [0.49, 1.99] s of the target presentation interval, feedback: [-1.51, -0.01] s relative to the end of the feedback interval). Significant p-values are highlighted in bold, and stars indicate that the differences remained significant after Bonferroni correction for multiple ($m = 3$) comparisons due to considering three time windows.

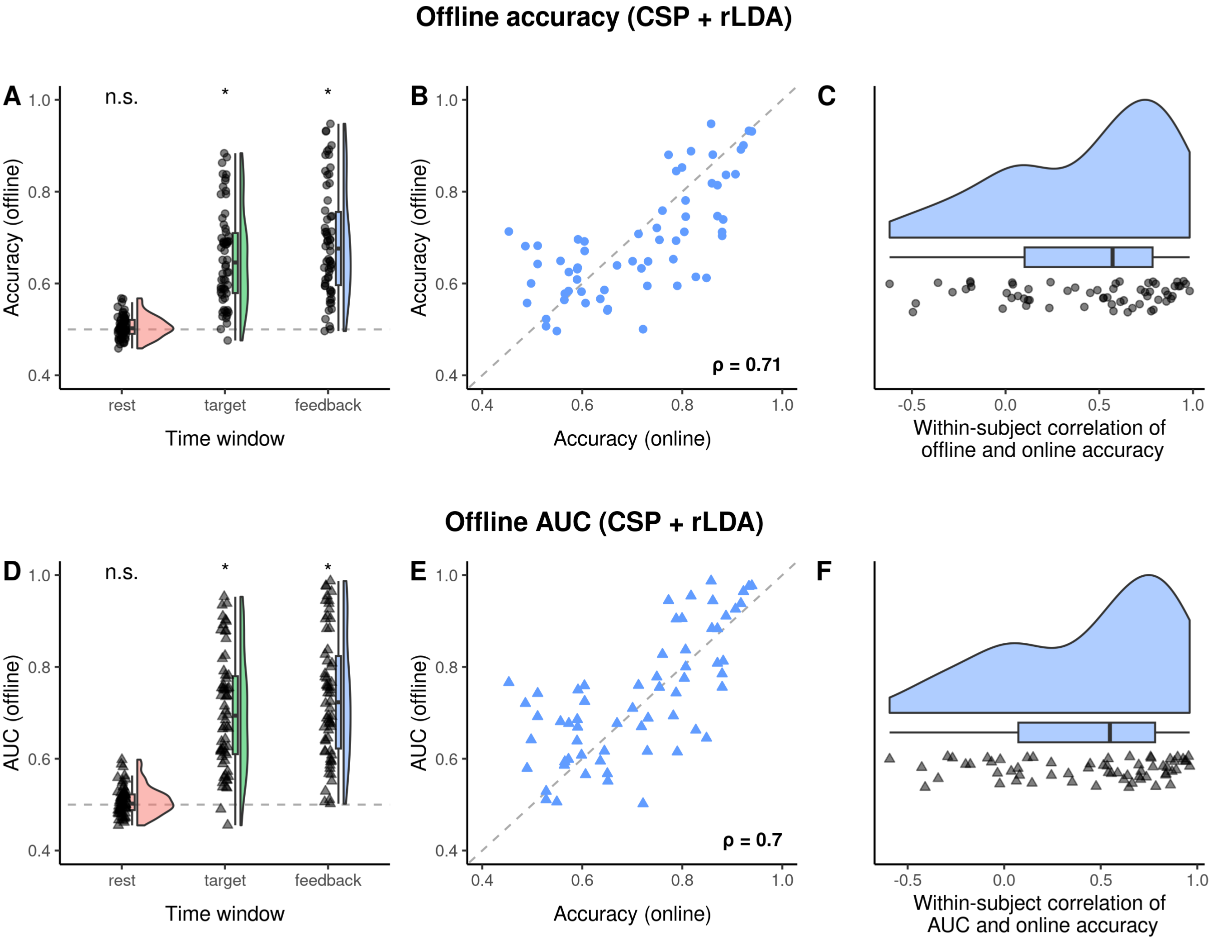


Figure S2: Values of offline accuracy and AUC obtained using CSP and regularized LDA in a 6-fold cross-validation. (A) Average accuracy for each subject in different time periods. No prediction during the inter-trial interval (rest) was possible, while both target presentation and feedback intervals allowed predicting the type of the imaginary movement. (B) Between-subject comparison of online and offline accuracy showed high correlation between the measures. Each point corresponds to a single subject. (C) Raincloud plot of within-subject correlation of online and offline accuracy. Each point corresponds to a single subject. (D-F) Same as (A-C) but for AUC instead of offline accuracy. Each triangle corresponds to a single subject.

### C. Laplacian-based analysis of phase synchronization

#### *C.1. Methods*

In order to make the Laplacian-based analysis of phase synchronization (PS) comparable to the source space one, we considered four channels (C3, C4, CP3, CP4), which are located directly over the pre- and postcentral gyri of both hemispheres. For CP3 and CP4, the surface Laplacian was applied by subtracting the average of neighboring channels CP1, CP5, C3, P3 and CP2, CP6, C4, P4, respectively. Values of PS were averaged over within-hemisphere (C3-CP3 and C4-CP4) and across-hemisphere (C3-C4, C3-CP4, CP3-C4, and CP3-CP4) connections. Values of SNR were averaged over all four channels and used in statistical analysis to correct for the effect of SNR on PS.

#### *C.2. Results*

The detailed statistical results are presented in Table S3. In particular, Laplace SNR had a significant effect on all PS measures except across-hemisphere LagCoh and coherence (Fig. S3). Among all PS measures, only within-hemisphere coherence had a significant positive effect on BCI performance after correction for SNR and multiple comparisons (Fig. S4). However, since neither ImCoh nor LagCoh showed a significant effect in the same analysis, the effect of coherence is likely to be driven by a zero-lag interaction and potentially by volume conduction. Finally, no evidence of longitudinal changes was observed for all PS measures before and after correction for SNR (Fig. S5). Overall, the observed results were very similar to the ones obtained with source space analysis.

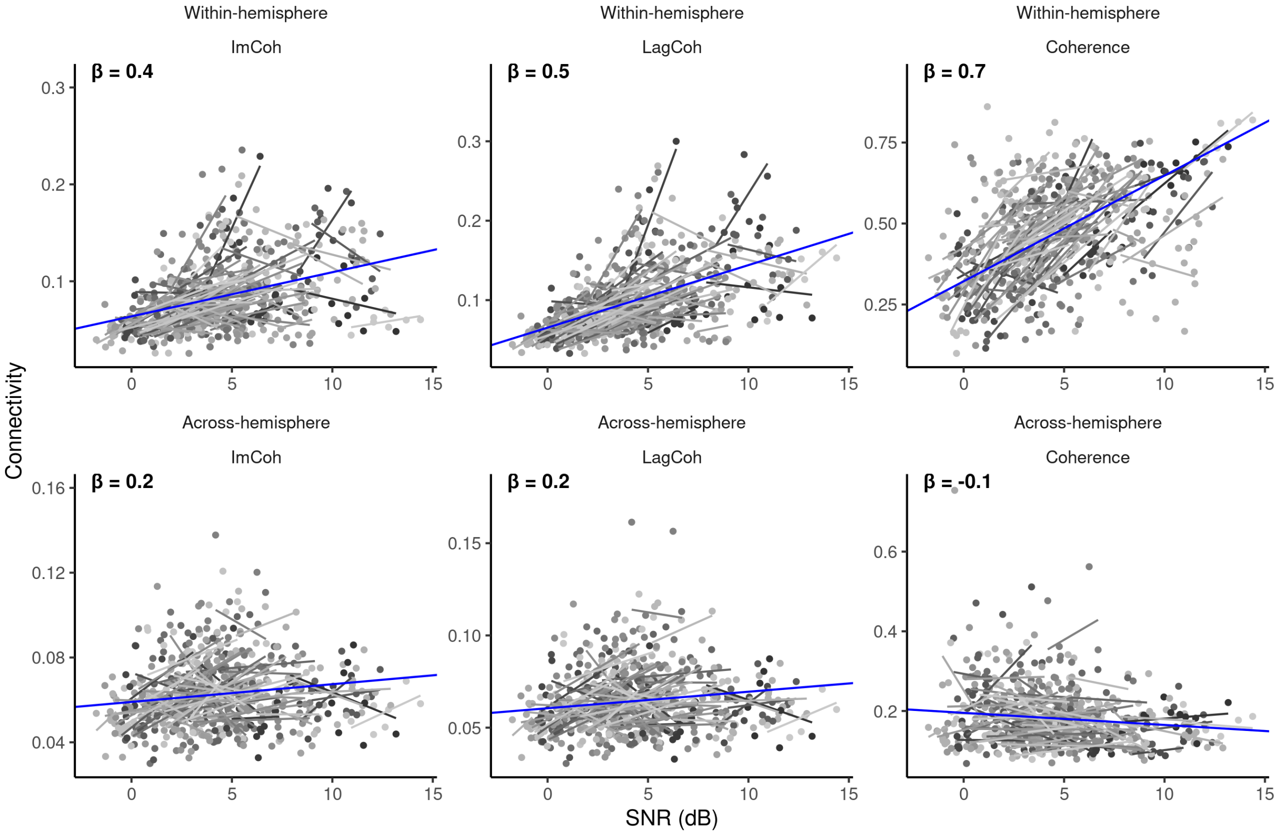


Figure S3: Laplace SNR had a significant effect on all measures of phase synchronization (PS) except across-hemisphere LagCoh and coherence.

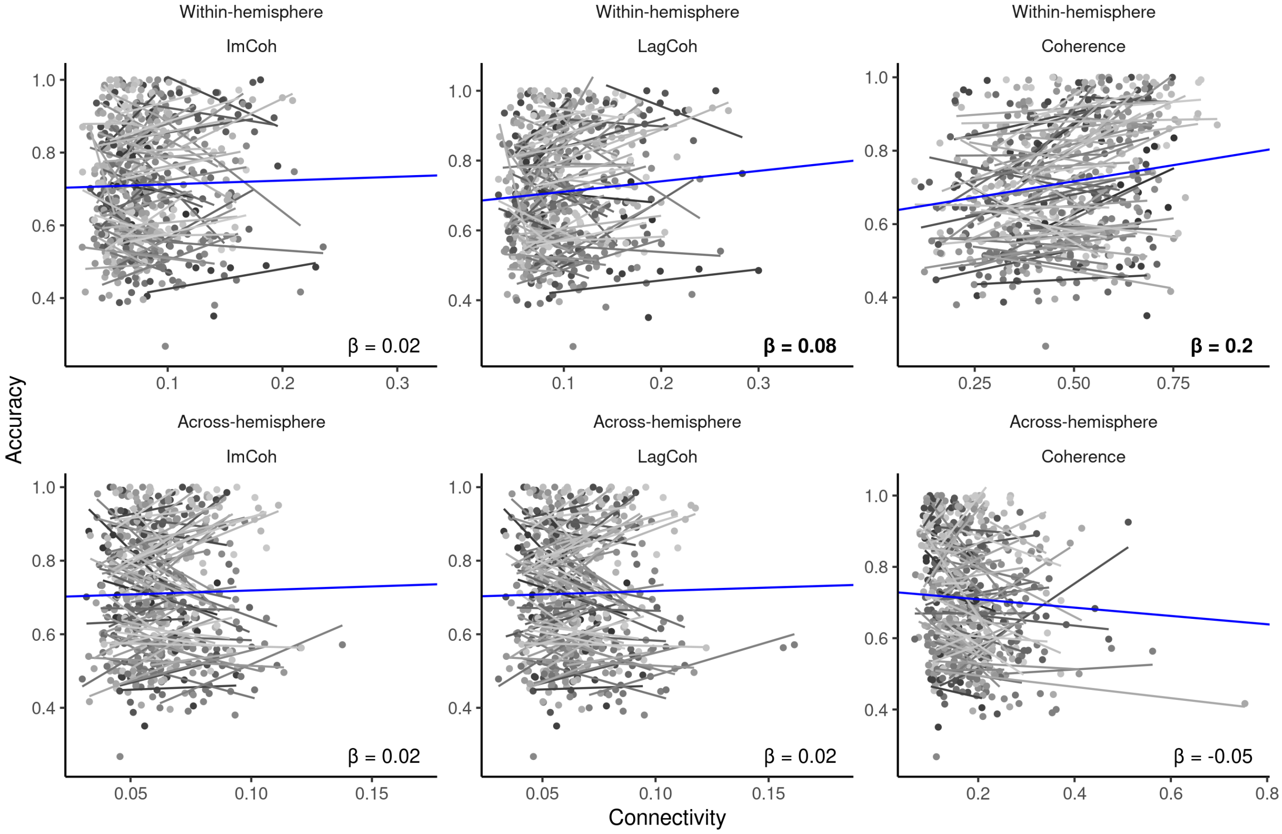


Figure S4: Among all PS measures, only within-hemisphere coherence had a significant positive effect on BCI performance after correction for SNR and multiple comparisons.

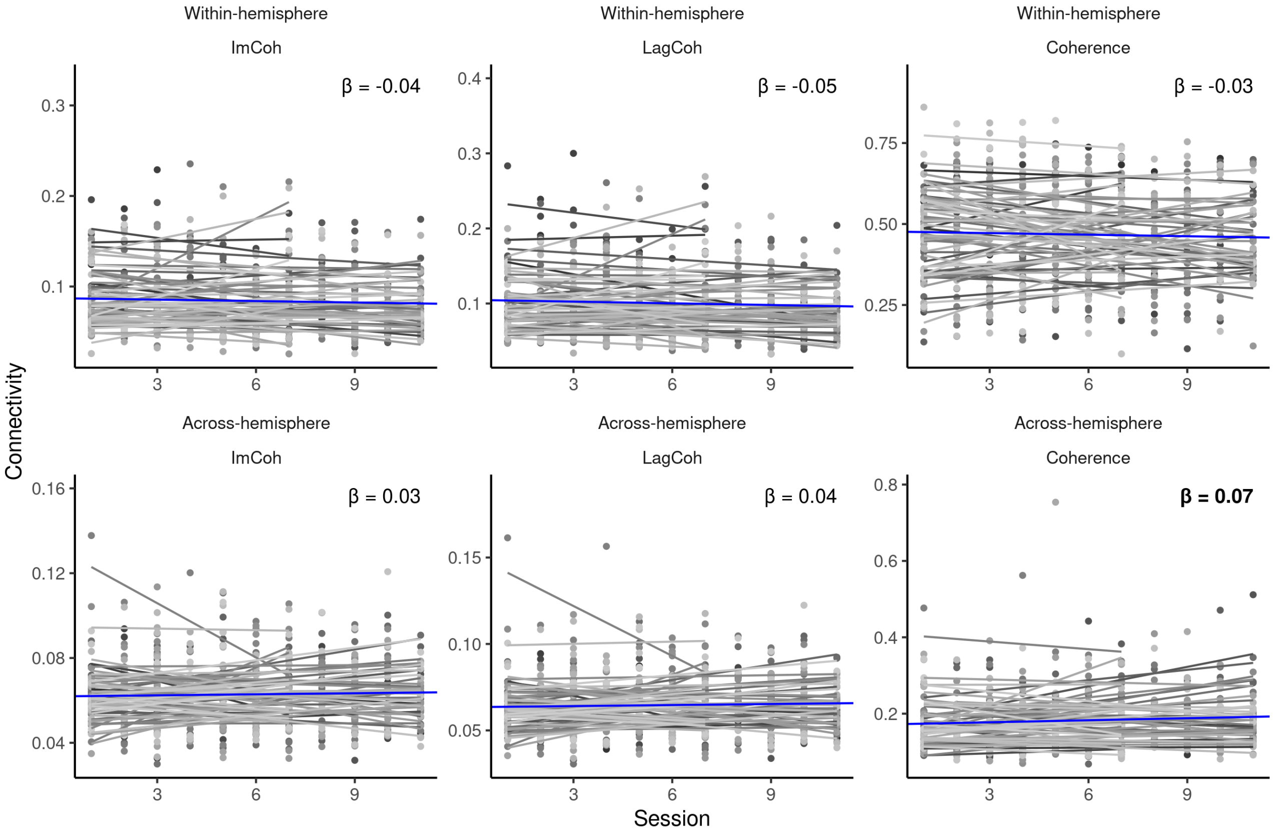


Figure S5: No evidence of longitudinal changes was observed for all measures of PS before and after correction for Laplace SNR.

| Predictor | Response | $\beta$ | t-value | p-value | 95% CI |
|---|---|---|---|---|---|
| SNR | WH ImCoh | 0.388 | 6.735 | <**0.001*** | $[0.275, 0.501]$ |
| SNR | WH LagCoh | 0.550 | 10.175 | <**0.001*** | $[0.443, 0.656]$ |
| SNR | WH Coherence | 0.669 | 12.200 | <**0.001*** | $[0.549, 0.788]$ |
| SNR | AH ImCoh | 0.160 | 2.696 | **0.008*** | $[0.041, 0.280]$ |
| SNR | AH LagCoh | 0.160 | 2.617 | **0.010** | $[0.037, 0.283]$ |
| SNR | AH Coherence | -0.126 | -2.086 | **0.038** | $[-0.245, -0.007]$ |
| WH ImCoh | Accuracy | 0.023 | 0.639 | 0.523 | $[-0.046, 0.092]$ |
| WH LagCoh | Accuracy | 0.079 | 2.174 | **0.030** | $[0.008, 0.150]$ |
| WH Coherence | Accuracy | 0.159 | 4.932 | <**0.001*** | $[0.096, 0.222]$ |
| WH ImCoh | Accuracy \| SNR | -0.027 | -0.762 | 0.446 | $[-0.097, 0.042]$ |
| WH LagCoh | Accuracy \| SNR | 0.007 | 0.180 | 0.857 | $[-0.068, 0.082]$ |
| WH Coherence | Accuracy \| SNR | 0.097 | 2.725 | **0.007*** | $[0.027, 0.168]$ |
| AH ImCoh | Accuracy | 0.021 | 0.686 | 0.493 | $[-0.039, 0.081]$ |
| AH LagCoh | Accuracy | 0.019 | 0.615 | 0.539 | $[-0.042, 0.081]$ |
| AH Coherence | Accuracy | -0.052 | -1.513 | 0.131 | $[-0.119, 0.015]$ |
| AH ImCoh | Accuracy \| SNR | -0.002 | -0.063 | 0.950 | $[-0.061, 0.057]$ |
| AH LagCoh | Accuracy \| SNR | -0.003 | -0.109 | 0.913 | $[-0.064, 0.057]$ |
| AH Coherence | Accuracy \| SNR | -0.040 | -1.196 | 0.232 | $[-0.106, 0.025]$ |
| Session | WH ImCoh | -0.044 | -1.381 | 0.168 | $[-0.108, 0.019]$ |
| Session | WH LagCoh | -0.052 | -1.680 | 0.094 | $[-0.113, 0.009]$ |
| Session | WH Coherence | -0.032 | -0.931 | 0.352 | $[-0.101, 0.036]$ |
| Session | WH ImCoh \| SNR | -0.034 | -1.094 | 0.274 | $[-0.096, 0.027]$ |
| Session | WH LagCoh \| SNR | -0.038 | -1.321 | 0.187 | $[-0.096, 0.019]$ |
| Session | WH Coherence \| SNR | -0.013 | -0.418 | 0.676 | $[-0.074, 0.048]$ |
| Session | AH ImCoh | 0.031 | 0.832 | 0.406 | $[-0.042, 0.105]$ |
| Session | AH LagCoh | 0.035 | 0.963 | 0.336 | $[-0.037, 0.107]$ |
| Session | AH Coherence | 0.073 | 2.204 | **0.028** | $[0.008, 0.138]$ |
| Session | AH ImCoh \| SNR | 0.036 | 0.979 | 0.328 | $[-0.037, 0.109]$ |
| Session | AH LagCoh \| SNR | 0.040 | 1.112 | 0.267 | $[-0.031, 0.111]$ |
| Session | AH Coherence \| SNR | 0.070 | 2.103 | **0.036** | $[0.005, 0.135]$ |

Table S3: Summary of the observed effects for the research questions in the Laplacian-based analysis of PS. Significant effects are highlighted in bold, and stars indicate that the effects remain significant after Bonferroni correction for multiple ($m = 6$) comparisons. WH and AH stand for within- and across-hemisphere values of PS metrics, respectively. Notation 'X | SNR' in the 'Response' column reflects that the effect of predictor on X was controlled for SNR.

### D. Additional tables and figures

| Sources | ROI according to the Harvard-Oxford atlas |
|---|---|
| **38** | **Left Precentral Gyrus** |
| **36** | **Right Precentral Gyrus** |
| **25** | **Right Postcentral Gyrus** |
| **24** | **Left Postcentral Gyrus** |
| 14 | Left Superior Parietal Lobule |
| 12 | Right Superior Parietal Lobule |
| 10 | Left Supramarginal Gyrus, posterior division |
| 8 | Right Supramarginal Gyrus, anterior division |
| 7 | Left Supramarginal Gyrus, anterior division |
| 6 | Right Middle Frontal Gyrus |
| 6 | Right Supramarginal Gyrus, posterior division |
| 5 | Left Middle Frontal Gyrus |
| 5 | Right Cingulate Gyrus, posterior division |
| 4 | Left Angular Gyrus |
| 4 | Left Precuneous Cortex |
| 4 | Right Central Opercular Cortex |
| 4 | Right Parietal Operculum Cortex |
| 3 | Left Cingulate Gyrus, posterior division |
| 3 | Right Juxtapositional Lobule Cortex (formerly Supplementary Motor Cortex) |
| 2 | Left Juxtapositional Lobule Cortex (formerly Supplementary Motor Cortex) |
| 1 | Left Lateral Occipital Cortex, superior division |
| 1 | Left Parietal Operculum Cortex |
| 1 | Right Superior Frontal Gyrus |
| 1 | Right Inferior Frontal Gyrus, pars opercularis |
| 1 | Right Lateral Occipital Cortex, superior division |
| 1 | Right Precuneous Cortex |

Table S4: Number of sources, which were considered task-relevant based on the source reconstruction of CSP patterns, for ROIs according to the Harvard-Oxford atlas. ROIs that were used in the analysis are highlighted in bold. ROIs that are not listed in the table did not contain any task-relevant sources.

| Predictor | Response | $\beta$ | t-value | p-value | 95% CI | Consistency |
|---|---|---|---|---|---|---|
| SNR | Accuracy | 0.266 | 7.879 | <**0.001*** | $[0.199, 0.333]$ | 1.00 |
| Session | SNR | 0.007 | 0.346 | 0.731 | $[-0.033, 0.046]$ | 1.00 |
| SNR | WH ImCoh | 0.195 | 17.731 | <**0.001*** | $[0.174, 0.217]$ | 0.92 |
| SNR | WH LagCoh | 0.503 | 40.446 | <**0.001*** | $[0.479, 0.528]$ | 1.00 |
| SNR | WH Coherence | 0.142 | 23.105 | <**0.001*** | $[0.129, 0.155]$ | 0.54 |
| SNR | AH ImCoh | 0.232 | 15.868 | <**0.001*** | $[0.203, 0.261]$ | 0.75 |
| SNR | AH LagCoh | 0.319 | 22.589 | <**0.001*** | $[0.292, 0.347]$ | 0.79 |
| SNR | AH Coherence | 0.198 | 16.351 | <**0.001*** | $[0.174, 0.223]$ | 0.33 |
| WH ImCoh | Accuracy | 0.023 | 4.190 | <**0.001*** | $[0.012, 0.034]$ | 0.42 |
| WH LagCoh | Accuracy | 0.055 | 8.845 | <**0.001*** | $[0.043, 0.068]$ | 0.54 |
| WH Coherence | Accuracy | 0.008 | 1.673 | 0.094 | $[-0.001, 0.018]$ | 0.71 |
| WH ImCoh | Accuracy \| SNR | 0.023 | 3.061 | **0.002*** | $[0.008, 0.038]$ | 0.00 |
| WH LagCoh | Accuracy \| SNR | 0.015 | 2.217 | **0.027** | $[0.002, 0.029]$ | 0.96 |
| WH Coherence | Accuracy \| SNR | -0.013 | -1.115 | 0.266 | $[-0.035, 0.010]$ | 1.00 |
| AH ImCoh | Accuracy | 0.033 | 5.775 | <**0.001*** | $[0.022, 0.044]$ | 0.00 |
| AH LagCoh | Accuracy | 0.034 | 5.857 | <**0.001*** | $[0.022, 0.045]$ | 0.00 |
| AH Coherence | Accuracy | 0.008 | 1.557 | 0.120 | $[-0.002, 0.018]$ | 1.00 |
| AH ImCoh | Accuracy \| SNR | 0.018 | 3.102 | **0.002*** | $[0.007, 0.029]$ | 0.00 |
| AH LagCoh | Accuracy \| SNR | 0.012 | 1.914 | 0.056 | $[0.000, 0.023]$ | 1.00 |
| AH Coherence | Accuracy \| SNR | -0.012 | -1.715 | 0.086 | $[-0.026, 0.002]$ | 1.00 |
| Session | WH ImCoh | -0.010 | -1.729 | 0.084 | $[-0.021, 0.001]$ | 0.96 |
| Session | WH LagCoh | -0.015 | -2.274 | **0.023** | $[-0.028, -0.002]$ | 1.00 |
| Session | WH Coherence | 0.002 | 0.606 | 0.545 | $[-0.004, 0.009]$ | 0.83 |
| Session | WH ImCoh \| SNR | -0.009 | -1.631 | 0.103 | $[-0.020, 0.002]$ | 0.96 |
| Session | WH LagCoh \| SNR | -0.013 | -2.122 | **0.034** | $[-0.026, -0.001]$ | 1.00 |
| Session | WH Coherence \| SNR | 0.003 | 0.829 | 0.407 | $[-0.004, 0.009]$ | 0.88 |
| Session | AH ImCoh | 0.011 | 1.513 | 0.130 | $[-0.003, 0.026]$ | 0.92 |
| Session | AH LagCoh | 0.018 | 2.536 | **0.011** | $[0.004, 0.033]$ | 0.92 |
| Session | AH Coherence | 0.026 | 4.152 | <**0.001*** | $[0.014, 0.038]$ | 0.29 |
| Session | AH ImCoh \| SNR | 0.012 | 1.641 | 0.101 | $[-0.002, 0.027]$ | 1.00 |
| Session | AH LagCoh \| SNR | 0.020 | 2.744 | **0.006*** | $[0.006, 0.034]$ | 0.08 |
| Session | AH Coherence \| SNR | 0.027 | 4.320 | <**0.001*** | $[0.014, 0.039]$ | 0.29 |

Table S5: Summary of the observed effects for the research questions in the joint multiverse analysis. Significant effects are highlighted in bold, and stars indicate that the effects remain significant after Bonferroni correction for multiple ($m = 6$) comparisons. WH and AH stand for within- and across-hemisphere values of PS metrics, respectively. Notation 'X | SNR' in the 'Response' column reflects that the effect of predictor on X was controlled for SNR.

| Interaction | Response | t-value | p-value |
|---|---|---|---|
| Band (NB) · Inverse (LCMV) | SNR | -12.43 | <**0.001***|
| | WH LagCoh | -4.90 | <**0.001*** |
| | WH Coherence | -2.44 | **0.04** |
| | AH ImCoh | -2.32 | **0.05** |
| | AH LagCoh | -9.00 | <**0.001*** |
| | AH Coherence | -5.45 | <**0.001*** |
| | WH ImCoh \| SNR | 2.50 | **0.03** |
| | WH LagCoh \| SNR | 3.01 | **0.01** |
| | AH LagCoh \| SNR | -2.37 | **0.04** |
| | AH Coherence \| SNR | -4.63 | <**0.001*** |
| Inverse (LCMV) · ROI Definition (Task-based) | WH LagCoh \| SNR | -2.77 | **0.02** |
| ROI Definition (Task-based) · ROI Method (3SVD) | WH ImCoh | 3.19 | **0.01** |
| | WH LagCoh | 2.58 | **0.03** |
| | WH Coherence | -2.31 | **0.05** |
| | WH ImCoh \| SNR | 3.15 | **0.01** |
| | WH LagCoh \| SNR | 4.24 | <**0.001*** |
| | WH Coherence \| SNR | -2.32 | **0.05** |
| ROI Definition (Task-based) · ROI Method (AVG-F) | WH ImCoh | -4.50 | <**0.001*** |
| | WH Coherence | 4.35 | <**0.001*** |
| | WH ImCoh \| SNR | -4.27 | <**0.001*** |
| | WH Coherence \| SNR | 4.51 | <**0.001*** |

Table S6: Interactions between processing steps that were significant for SNR or some of the PS measures (listed in the 'Response' column). Significant p-values are highlighted in bold, and stars indicate that the interaction remained significant after correction for multiple comparisons.

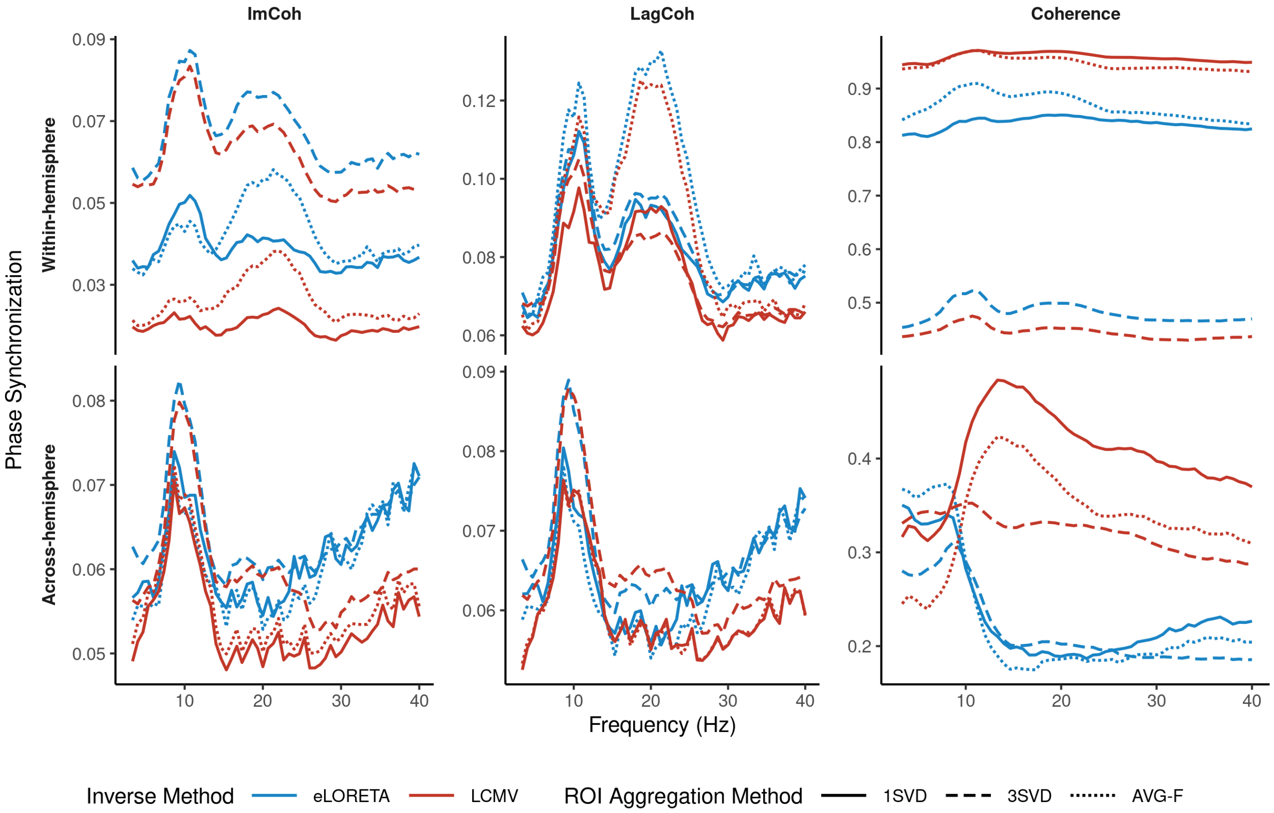


Figure S6: Grand average spectra of within- (top row) and across-hemisphere (bottom row) values of ImCoh, LagCoh, and coherence (columns: left to right) for the broadband pipelines with task-based definitions of ROIs and different ROI aggregation methods.

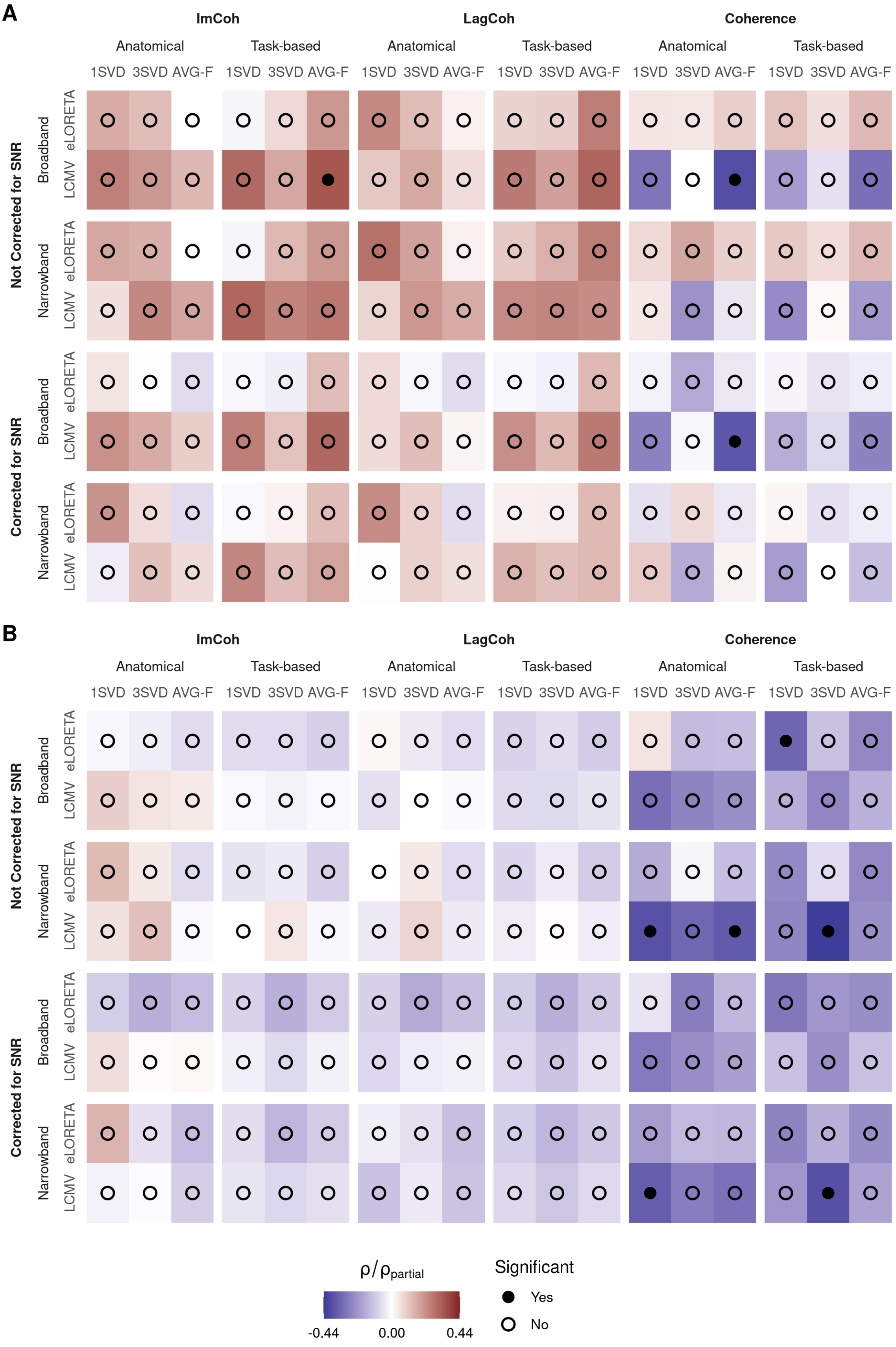


Figure S7: Between-subject effects of phase synchronization on performance as assessed with correlation ($\rho$, Not Corrected for SNR) and partial correlation ($\rho_{partial}$, Corrected for SNR). Panels (A) and (B) correspond to within- and across-hemisphere phase synchronization, respectively. Bonferroni correction for multiple ($m = 6$) comparisons was applied. Correlation values are coded with color.

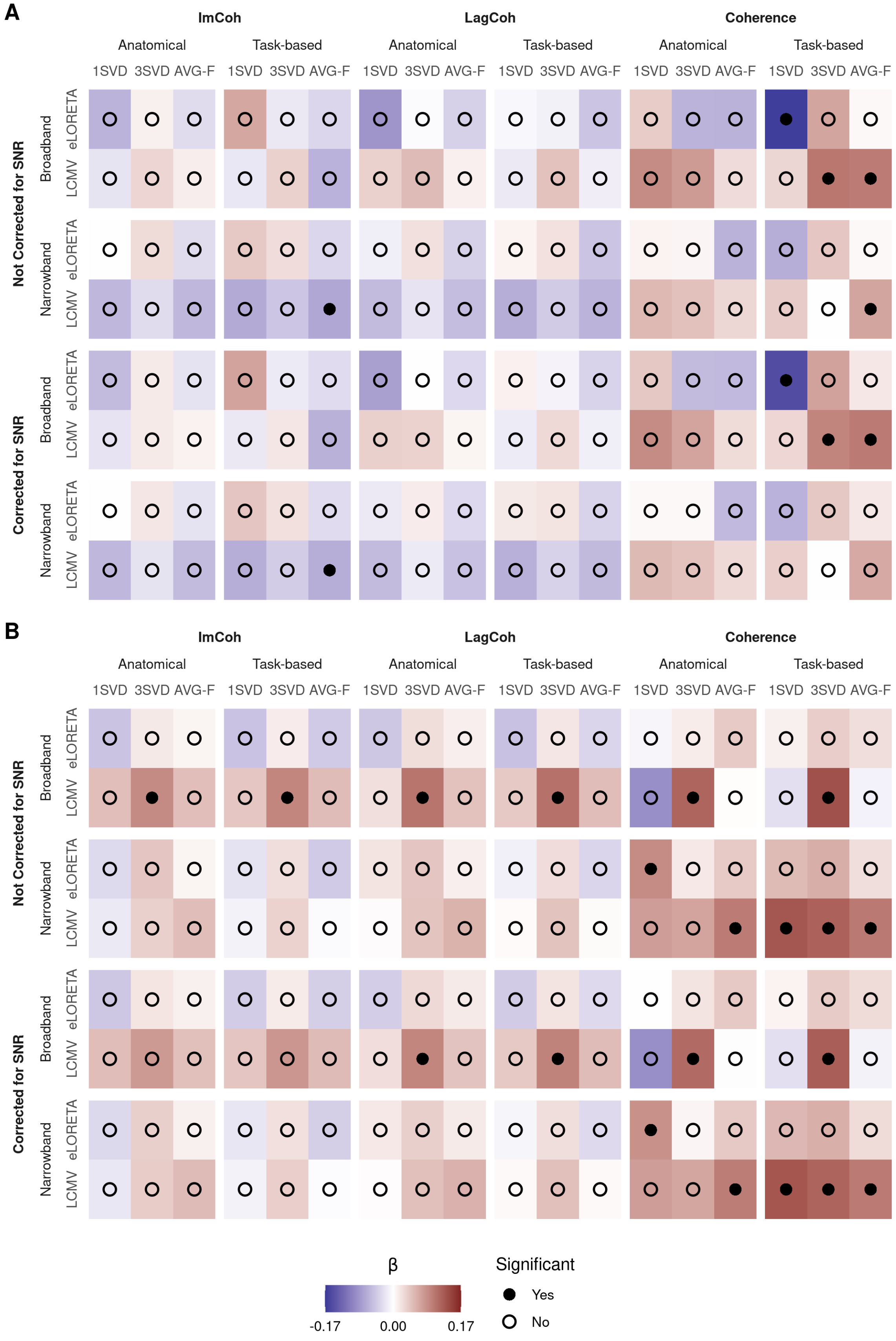


Figure S8: Longitudinal changes in phase synchronization throughout the BCI training. Panels (A) and (B) correspond to within- and across-hemisphere phase synchronization, respectively. Bonferroni correction for multiple ($m = 6$) comparisons was applied. Fixed effects of session on phase synchronization ($\beta$) are coded with color.

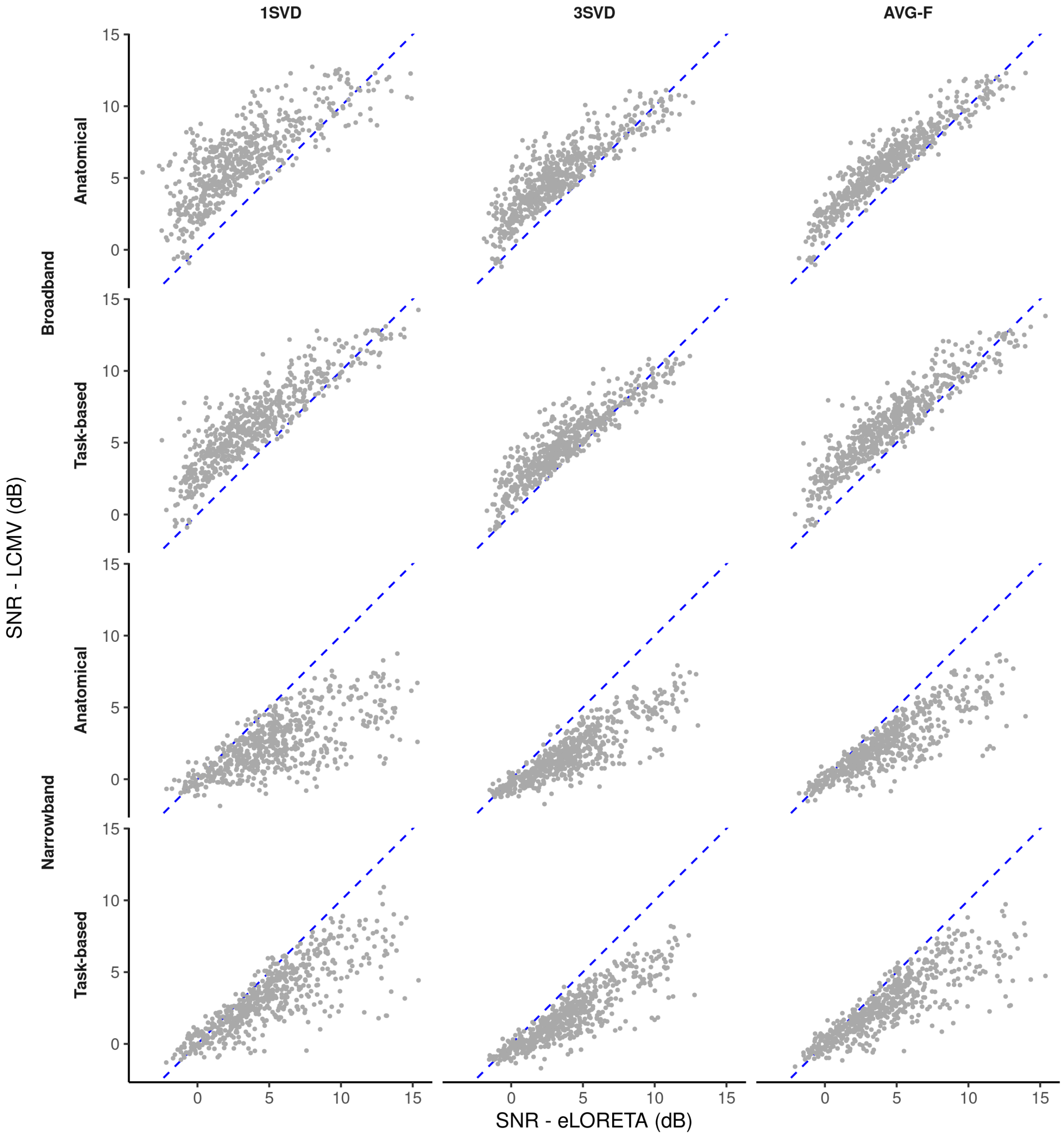


Figure S9: Interaction between filtering and the method for inverse modeling in the context of SNR. For broadband pipelines, LCMV led to higher values of SNR than eLORETA, and the difference was more pronounced for low values of SNR. For narrowband pipelines, SNR was higher when eLORETA was included in the pipeline. Blue dashed lines depict the area, where values of SNR estimated with LCMV and eLORETA are equal. For points above this line, SNR is higher when LCMV is used for inverse modeling, and vice versa. Each point corresponds to one training session.

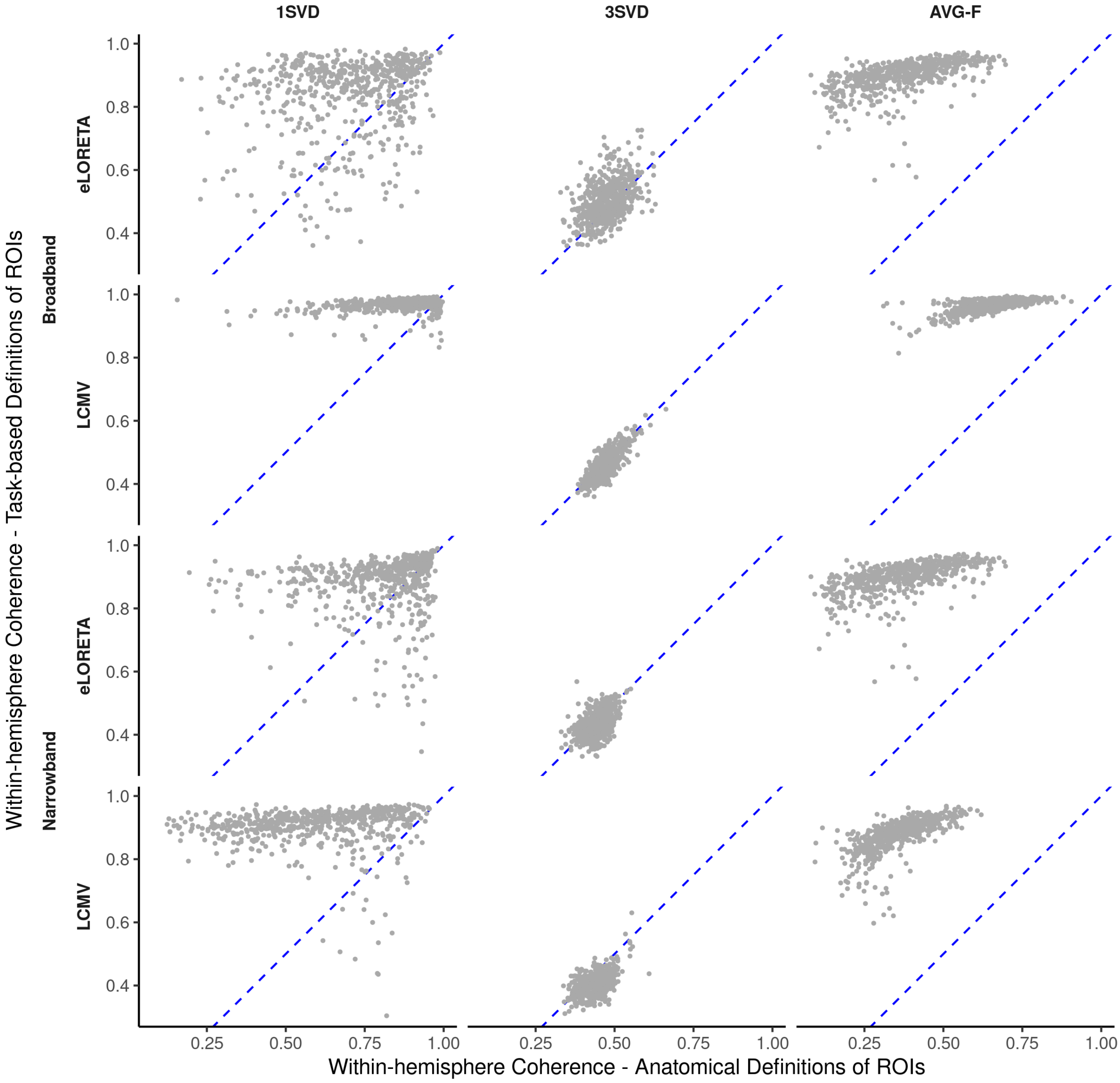


Figure S10: Interaction between the ROI definition and the method for extraction of ROI time series in the context of within-hemisphere coherence. When three SVD components per ROI were used, definition of the ROI had less impact on the coherence values compared to one SVD component or averaging with sign flip. Blue dashed lines depict the area, where values of coherence estimated with anatomical and task-based ROI definitions are equal. For points above this line, coherence is higher when task-based definitions are used, and vice versa. Each point corresponds to one training session.

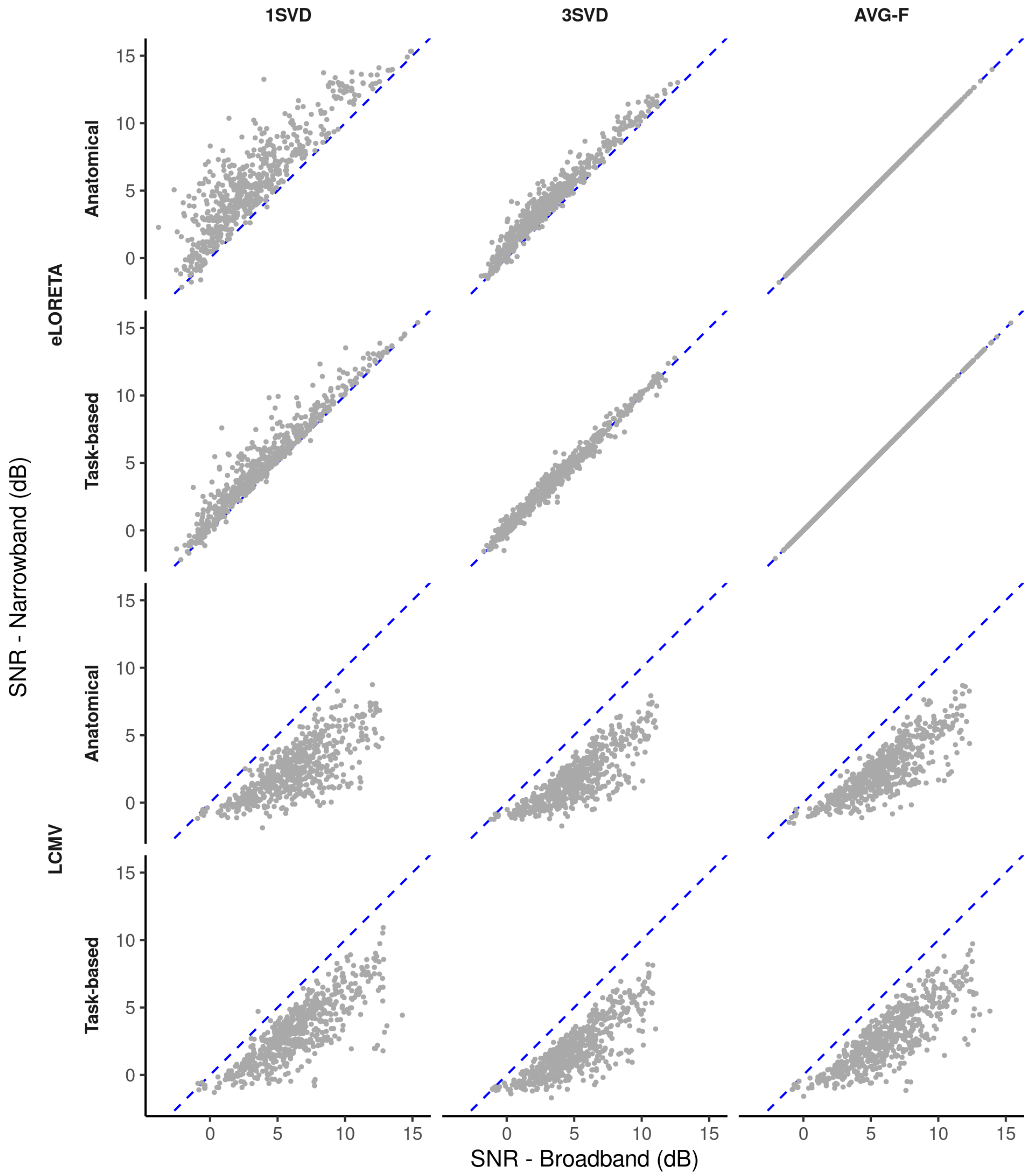


Figure S11: Interaction between filtering and the method for inverse modeling in the context of SNR. For pipelines that include eLORETA, SNR was higher when filtering in a narrow band was applied. At the same time, filtering decreased SNR for pipelines that include LCMV. Blue dashed lines depict the area, where values of SNR estimated with broadband and narrowband data are equal. For points above this line, SNR is higher when narrowband data is used, and vice versa. Each point corresponds to one training session.
